# Bacterial droplet-based single-cell RNA-seq reveals heterogeneity in bacterial populations and in response to antibiotic perturbation

**DOI:** 10.1101/2022.08.01.502326

**Authors:** Peijun Ma, Haley M. Amemiya, Lorrie L. He, Shivam J. Gandhi, Robert Nicol, Roby P. Bhattacharyya, Christopher S. Smillie, Deborah T. Hung

**Author notes:** Address correspondence to Deborah T. Hung.

## Abstract

We introduce BacDrop, a highly scalable technology for bacterial single-cell RNA sequencing that has overcome many challenges hindering the development of scRNA-seq in bacteria. BacDrop can be applied to thousands or millions of cells from both gram-negative and gram-positive species. It features universal ribosomal RNA depletion and combinatorial barcodes that enable multiplexing and massively parallel sequencing. We applied BacDrop to study *Klebsiella pneumoniae* clinical isolates and to elucidate their heterogeneous responses to antibiotic stress. In an unperturbed population presumed to be homogenous, we find within-population heterogeneity largely driven by the expression of mobile genetic elements that promote the evolution of antibiotic resistance. Under antibiotic perturbation, BacDrop revealed transcriptionally distinct subpopulations associated with different phenotypic outcomes including antibiotic persistence. BacDrop thus can capture cellular states that cannot be detected by bulk RNA-seq, which will unlock new microbiological insights into bacterial responses to perturbations and larger bacterial communities such as the microbiome.

## Introduction

Single-cell RNA-seq (scRNA-seq) has led to important discoveries in mammalian systems, resulting in a greater appreciation of the transcriptional heterogeneity of cell types and cell states [1–12]. While heterogeneity is essential for bacterial communities, bacterial scRNA-seq is still largely underdeveloped due to longstanding technical challenges. These include bacterial lysis, the absence of polyadenylated tails on messenger RNA (mRNA), and the paucity of mRNA molecules in a single bacterial cell [13], which collectively lead to degraded transcriptome coverage and quality relative to mammalian systems.

Recently, several bacterial scRNA-seq approaches have been described, including plate-based methods such as microSPLiT, PETRI-seq, and MATQ-seq [14–16], and probe-based methods such as par-seqFISH [17, 18].These plate-based scRNA-seq technologies enable genome-wide scRNA-seq, but they have to date been limited by the numbers of cells (scale) that can be studied. In addition, sequencing reads of these plate-based methods are dominated (>90%) by the overwhelmingly abundant rRNA, wasting a large portion of the sequencing investment. In contrast, the probe-based methods avoid the problem of abundant rRNA and have improved scale. However, these probe-based methods require prior knowledge of the genome(s) of interest to enable probe design and generation, with a pragmatic limitation to the numbers of genes that can be queried. Due to these limitations, studies using these previously reported methods have mainly focused on *between* population heterogeneity, with a focus on proof of principle, whereas *within* population heterogeneity, which requires the characterization of large number of cells from a single population at genome level, has not been extensively described.

One of the fundamental lessons from scRNA-seq of eukaryotic cells, especially of animals, is that because of the co-variation structure of gene expression within and between cells, profiling larger number of cells more shallowly is a more favorable experimental design than profiling a small number of cells deeply, and can better recover the statistical properties of cell populations and gene programs [7]. This is especially the case when molecular techniques limit the profile of each cell to a relatively low fraction of transcripts, randomly sampled from al transcripts in the cell. Combined with powerful algorithms, numerous scRNA-seq studies in eukaryotic systems have now demonstrated that novel cell types, states, dynamic trajectories, gene programs, and even features like spatial locations and cell interactions can be comprehensively recovered by such massively parallel methods, whereas per-cell coverage plays a less important role [1, 3–7, 9, 10, 19–26]. For instance, massively parallel single-cell and single-nucleus RNA-seq in human microglia detected novel subsets of cells and transformed our understanding of Alzheimer’s disease, even though only 1-2% of the entire genome (a few hundred genes) were detected per cell in these experiments [21–24]. Experimental and computational analyses have also demonstrated that low coverage and/or low sequencing depth is sufficient for effective cell clustering, detection of rare population, identification of biomarkers when analyzing a large number of cells, and conducting genetics “Perturb-Seq” screens [25–29]. The underlying principle of this observation is that scRNA-seq remains a sampling strategy, with only a portion of cells sampled from a population (scale) and a portion of RNA molecules sampled from a cell (coverage) [7]. Although higher coverage enables deep profiling of some cells and genes, it lacks the statistical power to recapitulate the phenotypic landscape of the population when the scale is limited. In contrast, because gene expression is structured due to shared regulatory mechanism, when a large number of cells are analyzed, even lower coverage of each cell allows the recovery of shared patterns such as clustering of similar cells to identify distinct subpopulations or ordering cells by Pseudotime to recover temporal trajectories [7, 26, 30]. Thus, to accurately capture the phenotypic landscape of the population and its statistical properties and distributions, rather than what is happening to one particular gene in any one particular cell, scale is more critical.

This same biological and statistical principle will be even more important in single cell transcriptional analysis of bacteria due to the inherent paucity of mRNA molecules in a single bacterial cell [13]. In addition, if applied to samples with high complexity or diversity, large numbers of cells are critical to ensure the capture of enough cells from any individual rare subpopulation or species, such as persister or heteroresistant subpopulations [13, 31–33] or the microbiome, mirroring the requirements in large-scale mammalian projects such as the Human Cell Atlas [2]. Therefore, methods for increasing bacterial scRNA-seq scale will be essential for studies in microbial systems.

Here we report the development and application of BacDrop, a droplet-based genome-wide massively parallel bacterial scRNA-seq technology. BacDrop has the flexibility to investigate a wide range of numbers of cells in one experiment, from thousands to millions of single bacterial cells, without requiring prior knowledge of the genome or probe-design. BacDrop also includes a universal and efficient rRNA-depletion step, reducing sequencing costs by at least ten-fold while simultaneously increasing information content. Furthermore, we demonstrated that BacDrop works on a variety of bacterial species, including the gram-negative *Escherichia coli*, *Klebsiella pneumoniae*, *Pseudomonas aeruginosa*, and the gram-positive bacterium *Enterococcus faecium*.

In this study, we applied BacDrop to assess any *within* population heterogeneity of *K. pneumoniae* and characterize its responses to antibiotics. *K. pneumoniae* is a gram-negative pathogen which has become one of the leading threats in the antibiotic resistance crisis, with increasing numbers of strains resistant to carbapenems, the last-resort antibiotic used to treat the most resistant infections [34]. Under stable and dynamic conditions (*i.e.,* without and with antibiotic perturbations), we identified important new features of heterogeneity that have heretofore been masked by bulk analysis in *K. pneumoniae* clinical isolates. In the absence of antibiotic perturbation, we found *within* population heterogeneity driven predominantly by mobile genetic elements (MGEs) which leads to high-level mutation frequencies promoting the evolution of antibiotic resistance, thus demonstrating novel subpopulation structure and function in a previously presumed homogenous population. After antibiotic perturbation, BacDrop revealed transcriptionally distinct subpopulations previously masked by bulk analysis that are associated with different phenotypic outcomes including decreased antibiotic efficacy and persister formation. With this demonstration of the power of BacDrop to illuminate heterogeneity both in static bacterial populations and their dynamic responses to perturbations, we propose that BacDrop has the potential to transform our understanding of bacterial survival, adaptation, and evolution, with numerous potential applications including the characterization of complex communities.

## Results

### BacDrop: a bacterial droplet-based massively parallel scRNA-seq technology

We developed BacDrop based on droplet-based microfluidics technology, which has the advantage of achieving much higher scale than plate-based technologies, leveraging the 10x Genomics^TM^ platform, a reliable and commercially available platform for scRNA-seq [9]. Two of the unique features of bacteria compared to mammalian cells are the challenge of lysing the bacterial cell wall, which requires harsher lysis conditions, while preserving RNA integrity, and the absence of mRNA polyadenylated tails, thus requiring an alternative approach to isolate mRNA (∼5%) from the vastly more abundant rRNA (∼95%). To overcome these problems, we adapted a previously reported cell fixation and permeabilization protocol to avoid cell lysis prior to droplet encapsulation [14, 16], and implemented universal rRNA and genomic DNA (gDNA) depletion steps within the permeabilized, fixed cells using RNase H and DNase I, respectively (Fig. 1A, Fig. S1A).

**Figure 1.**
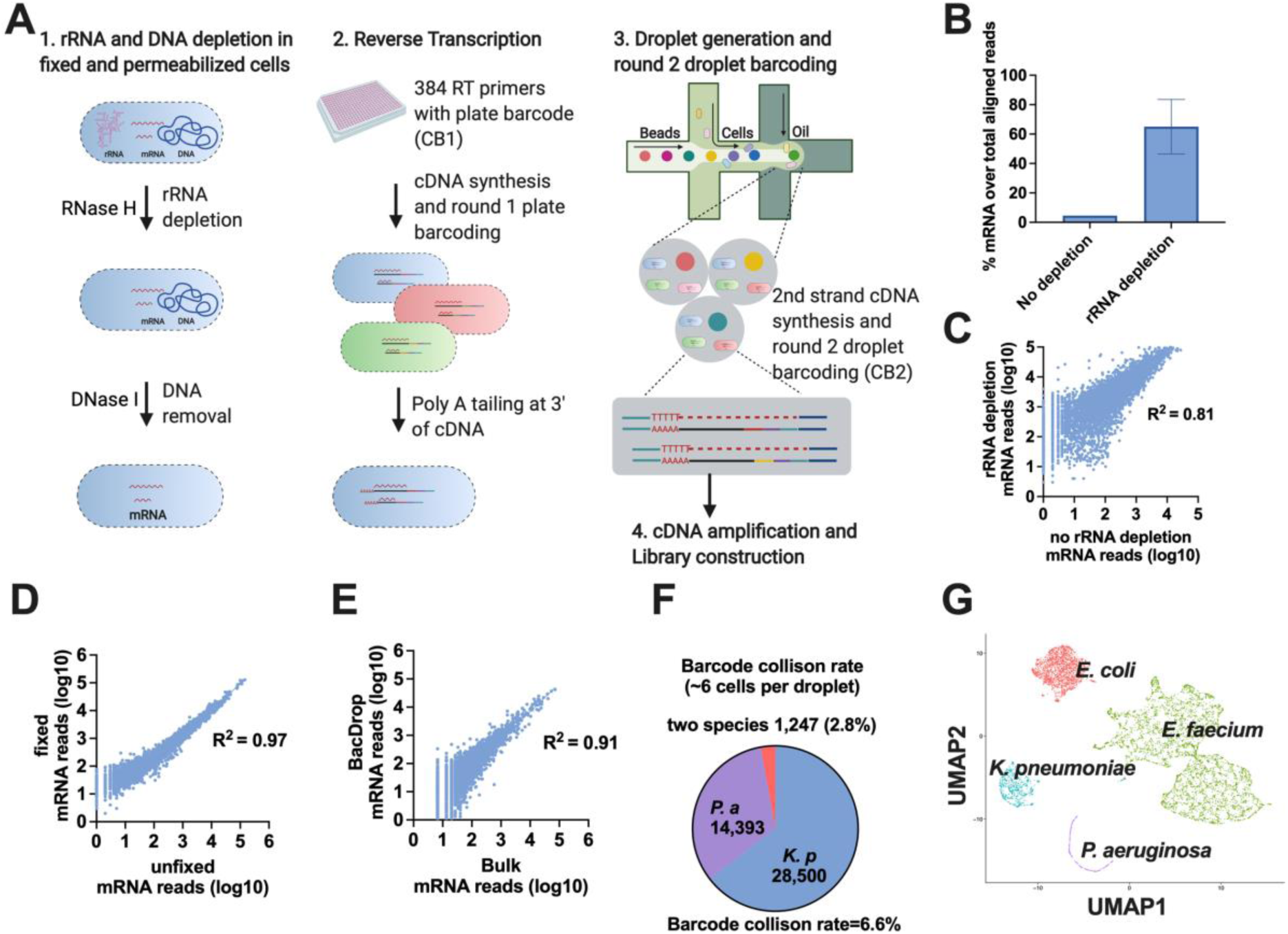
BacDrop: a bacterial droplet-based massively parallel scRNA-seq technology. (A) BacDrop workflow. Following cell fixation and permeabilization, rRNA and gDNA is depleted from cells in bulk. Then CB1 and UMIs are added to the 5’ end of cDNA via RT reactions (round 1 plate barcoding) in pools of ∼200,000 cells per well in 96- or 384-well plates. After round 1 plate barcoding, all cells are pooled and cDNA is polyadenylated at the 3’ end using terminal transferase, followed by droplet generation and round 2 droplet barcoding. The 3’ poly-A tail of cDNA enables second strand synthesis using oligo-dT primers. Round 2 droplet barcoding is achieved via second strand cDNA synthesis and 4 capturing cycles by barcoded primers (on 10x gel beads) in droplets. The successfully captured cDNA contains UMIs, CB1 and CB2, as well as adaptor sequences at both 5’ and 3’ end. Each cell is identified by a combination of CB1 and CB2. (B) rRNA depletion inside fixed and permeabilized cells significantly increases the percentage of total reads that align to mRNA in BacDrop libraries of *K. pneumoniae*. (C) rRNA depletion does not affect mRNA profiles. BacDrop libraries were constructed using in *K. pneumoniae* samples with or without rRNA depletion, and a linear regression model was fitted to the mRNA counts from each library. (D) Cell fixation and permeabilization does not affect mRNA profiles. Bulk RNA-seq results derived from Trizol-extracted RNA samples of *K. pneumoniae* versus RNA derived from fixed and permeabilized *K. pneumoniae* cells were highly correlated. (E) Single-cell BacDrop results are highly correlated with bulk RNA-seq results when analyzed in bulk mode (without cell barcode extraction). (F) BacDrop has low barcode collision rates in an experiment where 2 million bacterial cells, mixed with *K. pneumoniae* and *P. aeruginosa* cells, were loaded into one 10x channel (∼6 cells per droplet). About 2.8 % of the cells was assigned to two species, resulting in a 6.6% barcode collision rate. (G) BacDrop was performed on 4 different bacterial species including the gram-negative *E. coli*, *K. pneumoniae*, *P. aeruginosa*, and the gram-positive bacterium *E. faecium*. Uniform Manifold Approximation and Projection (UMAP) of this mixed population shows the separation of different species, colored by species identity.

Droplet-based single cell approaches require Poisson loading at mean droplet occupancy values (λ) far less than 1 to ensure that multiple cells are not encapsulated within a single droplet [1] and thus do not share the same barcode. Because of this low occupancy value, Poisson loading limits numbers of cells that can be loaded in each microfluidic channel. To avoid this problem, we utilized a multi-step barcoding strategy wherein cells are uniquely identified using the combination of two barcodes: a “plate barcode” (CB1) as a pre-indexing step corresponding to one of a possible 384 different barcodes, and a second “droplet barcode” (CB2) that uniquely identifies each droplet, together constituting the cell barcode. A unique molecular identifier (UMI) is also included as part of CB1 to enable identification of unique transcripts via removal of PCR duplicates. This pre-indexing strategy has recently been successfully applied to mammalian droplet-based scRNA-seq [35].

The pre-indexing with CB1 was achieved via reverse transcription (RT) reactions using random hexamer priming in a 384-well plate (plate barcoding), with ∼200,000 cells in each well containing one of 384 uniquely barcoded RT primers[35] (Fig. 1A, Table S1). Template switching is less efficient for second strand cDNA synthesis from bacterial than mammalian mRNA due to the lack of proper modifications at the 5’ end of bacterial mRNA. We thus used Terminal Transferase (TdT) to append poly-A tails to the 3’ end of cDNA to facilitate second strand cDNA synthesis inside droplets. By pre-indexing, we were able to load each droplet with multiple cells, increasing the scale significantly from a λ of ∼0.3 for conventional droplet scRNA-seq experiments to a λ of >1 for our approach with multiple cells in each drop.

For droplet generation and round 2 droplet barcoding (CB2), we utilized the commercially available single-cell kit from 10x Genomics^TM^ (see Methods) and loaded 3-6 cells per droplet. After round 2 droplet barcoding, cDNA from each cell was now labeled with a unique cell barcode, a combination of CB1 and CB2. cDNA libraries were then constructed using a protocol modified from the Illumina Nextera XT DNA library construction kit (Fig. 1A, Fig. S1B).

### Determining the technical performance of BacDrop

To characterize the performance of BacDrop, we tested the efficiency of rRNA depletion, the effect of fixation on gene expression profiles, the congruence of transcriptional profiles with bulk RNA-seq libraries, barcode collision rates, and generalizability to different bacterial species. After rRNA depletion, the fraction of mRNA reads increased from ∼5% to 50-90% of the total aligned reads, while preserving mRNA profiles relative to non-depleted samples (R^2^ = 0.81; Fig. 1B-1C; Fig. S1C). In addition, we confirmed that mRNA profiles were not affected by cell fixation in bulk experiments (R^2^ = 0.97; Fig. 1d), and that BacDrop produces transcriptional profiles that are well-correlated with those generated by the traditional bulk RNA-seq method (R^2^ = 0.91; Fig. 1D-1E).

We confirmed that the two-step barcoding strategy, which was used to enable loading of multiple cells per droplet, resulted in a minimal number of cells labelled with the same barcodes, that is the cell barcode collision rates after two rounds of barcoding. We conducted round 1 plate barcoding using 384 RT primers and round 2 droplet barcoding containing on average 6 cells per droplet, though with a range of cell numbers per droplet, on a mixed population of two distinguishable species, *K. pneumoniae* and *P. aeruginosa*. Of 44,140 cells that passed quality control at a sequencing depth of ∼1,000 reads per cell, 42,893 cells (97.2%) had > 99% of UMIs aligning to a single species, whereas the other 1,247 cells (2.8%) had > 1% of UMIs aligning to both species (Fig. 1F). The resulting barcode collision rate of 6.6% compares to published methods for eukaryotic droplet scRNAseq [1, 35]. We performed all subsequent experiments with < 3 cells per droplet, which would yield a library containing maximally a million cells in each 10x channel with an even lower collision rate.

To demonstrate the generalizability of BacDrop to different bacterial species and the ability of BacDrop to differentiate among multiple species, we applied BacDrop to the gram-negative *E. coli*, *K. pneumoniae*, *P. aeruginosa*, and the gram-positive *E. faecium* (Fig. 1G, Table S2). We performed cell fixation, permeabilization, rRNA and gDNA depletion, and round 1 plate barcoding separately on individual cultures of these four species, then mixed all four strains together into a single pool containing roughly one million cells for round 2 droplet barcoding and library construction. Each strain was thus labeled with a unique set of CB1 (Table S1), allowing us to track and validate the accuracy of species identification in downstream analyses. We used the standard workflow of Seurat 3 for cell clustering and subpopulation identification[36]. All four species were clearly distinguished in the analysis and the RNase-H based rRNA depletion worked efficiently across all four species (Fig. 1G, Fig. S1D-1E). Cells from *E. faecium* fell into several subpopulations distinguished by differential expression of two highly expressed housekeeping genes (*ef-tu* and *ef-g*; log2 fold change ∼ 0.6). Of note, the variation of cell numbers among these four species in the sequenced library was due to cell loss before round 2 droplet barcoding. Specifically, this was due to the loss of a larger fraction of *P. aeruginosa* cells, which are smaller and thus not recovered as efficiently during the centrifugation steps (Fig. S1F). For future experiments, we would only pool samples right before round 2 droplet barcoding to ensure equal number of cells from each sample are loaded into the 10x channel for cell barcoding and library construction or use centrifugation speeds that have been optimized for each species or sample.

We next examined the transcriptome coverage of BacDrop in *E. coli* and *K. pneumoniae*. We generated two small libraries containing either ∼10,000 cells of *E. coli* or ∼12,000 cells of *K. pneumoniae* (Fig. S1G-1H). Both libraries were sequenced with ∼80,000 reads per cell, and we recovered roughly 4,000 or 6,000 cells, containing at least 15 mRNA genes per cell, respectively. We detected the expression of ∼70% genes of the *E. coli* genome and ∼80% genes of the *K. pneumoniae* genome when we analyzed all cells together (Table S3, Table S4). At the single-cell level, we detected an average of 90 and 88 mRNA genes per cell, respectively, which is comparable to other reported bacterial scRNA-seq methods [14, 16]. We also generated a large library from ∼1 million *K. pneumoniae* cells and sequenced with ∼5,000 reads per cell. We recovered ∼60,000 cells with at least 15 mRNA genes per cell and detected the expression of 96% genes of the entire genome when we analyzed all cells together (Table S5). At the single-cell level we detected an average of 30 mRNA genes per cell across all 60,000 cells. However, from this large library, the top 3,000 high-quality cells had an average of 127 mRNA genes detected per cell (Fig. S1I). Since this large library was only sequenced with ∼5,000 reads per cell, we expect to detect a higher number of mRNA genes per cell with increased sequencing depth, as illustrated with the smaller libraries.

Finally, we assessed BacDrop’s sensitivity to distinguish different expression levels of a gene. We created a heterogeneous population containing three *E. coli* strains constitutively expressing *gfp* at different levels (Table S2). We used flow cytometry to confirm the differing expression levels of *gfp* in these strains and estimated the mRNA copy numbers of *gfp* in each of these strains using RT-qPCR [13] (Fig. 2A-2B). The estimated mRNA copy numbers per cell were 1-5 for the gfp.low strain, 9-30 for the gfp.mid strain, and 30-70 for the gfp.high strain. We barcoded each strain with a unique set of CB1 and mixed ∼3,300 cells from each strain for round 2 droplet barcoding to generate a BacDrop library. We found BacDrop can distinguish cells based on their *gfp* expression levels. Importantly, the *gfp* expression levels detected via BacDrop were consistent with the flow cytometry and RT-qPCR results (Fig. 2C-2D), suggesting that BacDrop can reliably identify subpopulations with differentially expressed genes.

**Figure 2.**
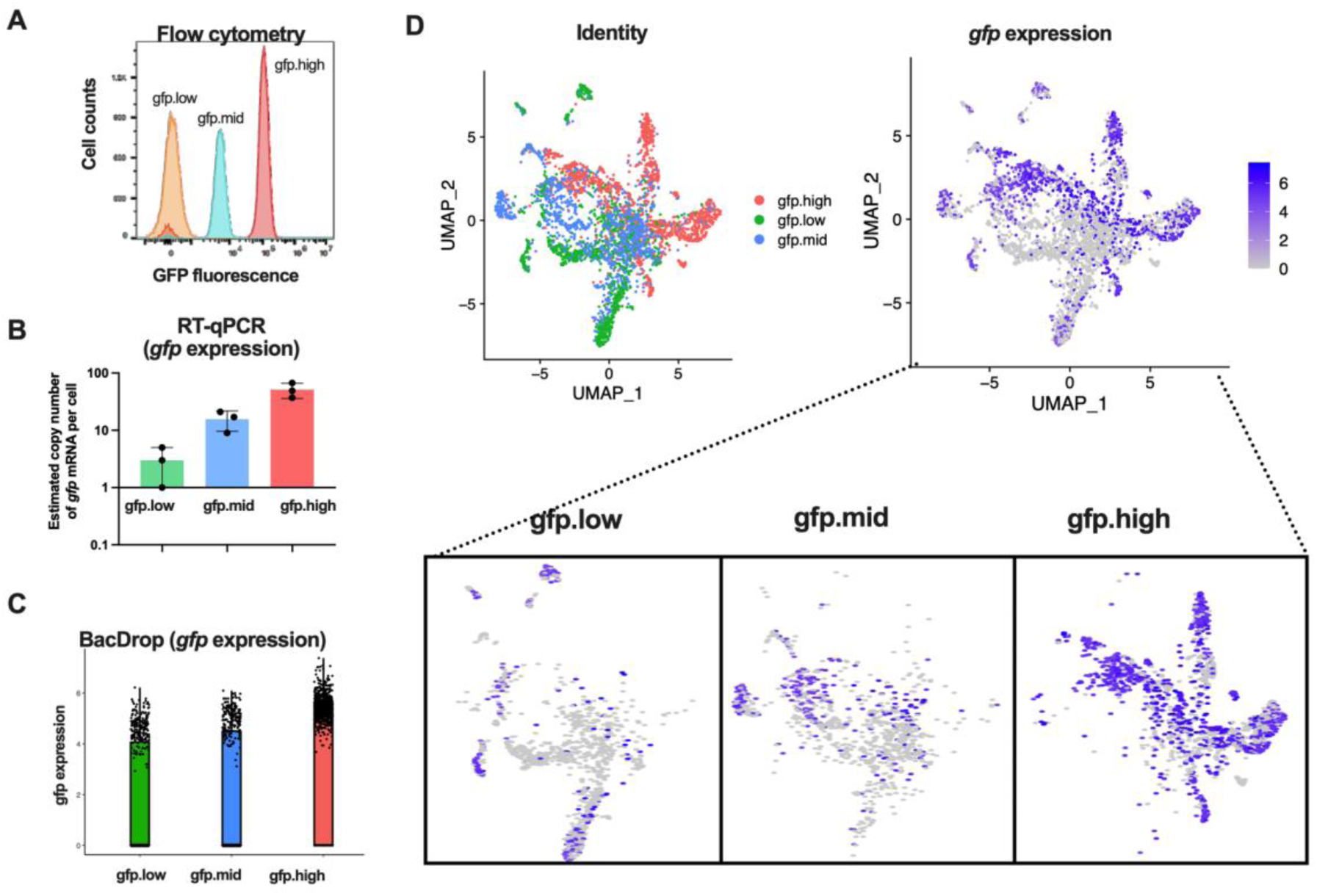
BacDrop has good sensitivity to distinguish cells based on the expression level of *gfp*. (A-B) The expression levels of *gfp* in three *E. coli* strains were confirmed using flow cytometry. (A) and the mRNA copy numbers of *gfp* in the sorted population were estimated using RT-qPCR. (B) The mRNA copy numbers were 1-5 for the gfp.low strain, 9-30 for the gfp.mid strain, and 30-70 for the gfp.high strain. Roughly 3,300 cells from each strain were mixed to create a heterogeneous population and a BacDrop library was generated. (C) BacDrop results of *gfp* expression in three strains. (D) Identify of cells and their *gfp* expression levels.

Together, these results suggest that BacDrop is a robust and reliable technology with sufficient coverage and sensitivity that can be applied across different numbers of cells and species. We thus proceeded to test its capability in revealing biological heterogeneity in a single *K. pneumoniae* clinical isolate, using these experiments to additionally demonstrate BacDrop’s reproducibility.

### Validating BacDrop’s ability to identify subpopulations of a single bacterial isolate

To validate that BacDrop could reproducibly identify subpopulations of cells of the same isolate, we applied BacDrop to the antibiotic-susceptible clinical isolate, *K. pneumoniae* MGH66 in the absence and presence of antibiotic perturbations (Fig. 3A, Table S2). We performed two biological replicates, denoted as Replicate 1 and Replicate 2. For each replicate, we split a MGH66 culture (OD_600_ ∼0.2) into four identical cultures, then treated three of them with antibiotics with different mechanisms of action, including inhibition of cell-wall synthesis (meropenem), DNA synthesis (ciprofloxacin), and protein synthesis (gentamicin). The fourth culture was left untreated (see Methods). Bulk RNA-seq on samples collected using the same treatment schemes revealed distinct cellular responses to each antibiotic relative to the untreated control. As expected (Fig. 3B), ciprofloxacin induced genes involved in the SOS response, *e.g*., *recA*, while gentamicin treatment induced a group of heat shock chaperone proteins, *e.g., ibpB*. In contrast, at 30 minutes, minimal transcriptional responses were observed in bulk from meropenem treatment, under the condition used (OD_600_ ∼ 0.2, meropenem concentration 2 μg/mL), which is consistent with previous observations [37].

**Figure 3.**
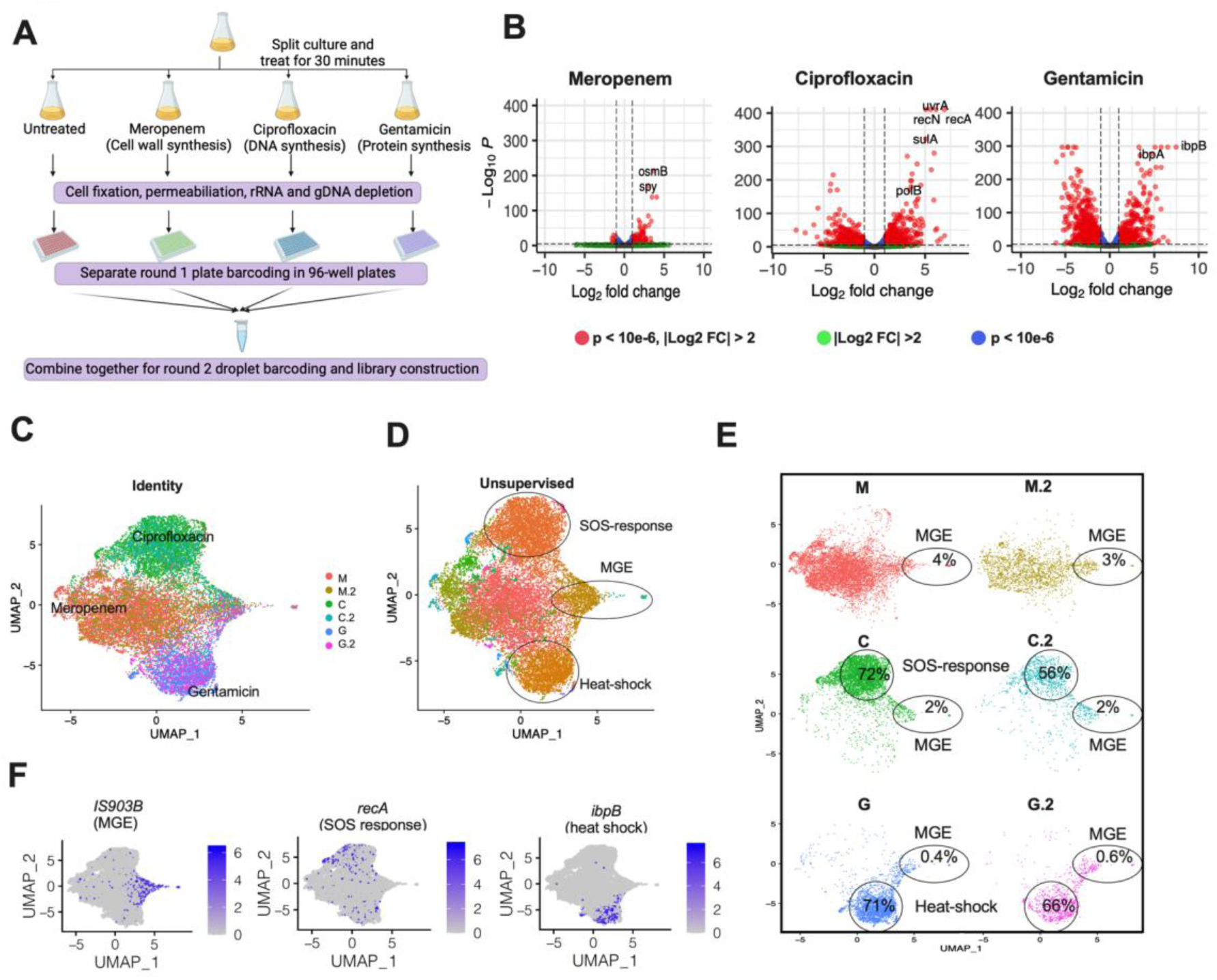
Validating BacDrop’s ability to distinguish subpopulations based on distinct responses to different antibiotic treatments. (A) Creation of a BacDrop library containing cells of the same bacterial strain under 4 different conditions. Cultures of MGH66 were treated for 30 minutes with antibiotics: meropenem, 2 µg/mL; ciprofloxacin, 2.5 µg/mL; gentamicin, 4 µg/mL; and no antibiotic control. Cells were collected and processed separately until after round 1 plate barcoding. The four samples were then pooled for round 2 droplet barcoding and library construction. Two biological replicates were performed. (B) Bulk RNA-seq results of cells exposed to the same antibiotic conditions as in (A). The abundance for each treated condition and comparison between the treated and untreated cultures are shown as well as significantly up- and down- regulated genes from each treatment (performed in triplicates). (C) UMAP plotted based on the original identity of the 6 samples treated with meropenem (M and its replicate M.2), ciprofloxacin (C and its replicate C.2), and gentamicin (G and its replicate G.2). (D) Unsupervised UMAP showed three clusters with significantly (p < 0.05) higher expression of genes in the SOS-response pathway, heat-shock response, and genes encoding an IS903B transposase (MGE). (E) No strong batch effect was observed between the two biological replicates with the same treatment conditions. 56%-72% of the ciprofloxacin-treated cells were clustered in the SOS-response cluster, and 66%-71% of the gentamicin-treated cells were clustered in the heat-shock response cluster. All 6 samples contained cells highly expressing *IS903B* (MGE cluster). (F) Expression of a representative gene from each cluster was highlighted on the UMAP.

To validate BacDrop, we applied round 1 plate barcoding to each of these 4 samples separately using one of 4 distinct sets of CB1 (96 different CB1s corresponded to each sample, Table S1). The distinct CB1 set identities allowed us to confirm the original identity of these 4 samples and associate them with the corresponding antibiotic exposure. We then mixed all cells from the different treatments for round 2 droplet barcoding and library construction (Fig. 3A). Each of the replicate libraries contained roughly one million cells. Replicate 1 was sequenced with ∼5 billion paired-end reads (∼5,000 reads per cell) and Replicate 2 with ∼3 billion paired-end reads (∼3,000 reads per cell).

To confirm BacDrop’s robustness and reproducibility, we compared the two replicate experiments, each involving 4 samples (3 antibiotic treated and 1 untreated). For the depth of sequencing performed for each experiment, we obtained ∼80,000 cells with ≥15 mRNA genes per cell, with an average of ∼30 unique mRNA genes detected per cell. No strong batch effect was observed between the replicates. Treatment with the different antibiotics, ciprofloxacin, gentamicin, or meropenem, resulted in cells clustering based on their treatment conditions, (Fig. 3C-3F). There was some overlap between the meropenem-treated and untreated samples (Fig. S2), consistent with their bulk RNA-seq results which suggest relatively minimal transcriptional response to meropenem, at least on the population level (Fig. 3B). Across both replicates, 56%- 72% of the ciprofloxacin-treated cells belonged to the SOS-response cluster, while 66%-71% of the gentamicin-treated cells belonged to the heat-shock response cluster, with good reproducibility across the two replicates (Fig. 3E). These experiments confirmed that BacDrop could effectively and reproducibly identify population heterogeneity.

### BacDrop reveals *within* population heterogeneity with subpopulations driven largely by the expression of MGEs

We next analyzed in depth the untreated culture of MGH66 at the single cell level to understand if it contained any previously unrecognized *within* population heterogeneity in transcriptional states. From this untreated condition we recovered ∼50,000 cells with at least 15 unique mRNA genes detected in each cell, and we identified two major subpopulations in both replicates using an unsupervised clustering approach (see Methods; Fig. 4A, Fig. S3). While the majority of cells fell into one major homogenous subpopulation, 2,191 cells (∼4.5%) fell into the MGE subpopulation driven by IS903B (Fig. 4B), which has 83 copies in MGH66 genome. (In fact, this MGE subpopulation is present in all 8 samples, untreated and antibiotic treated replicates, suggesting both that the presence of this MGE population is a robust phenomenon and that BacDrop is reproducible (Fig. 3D-3F, 4A).)

**Figure 4.**
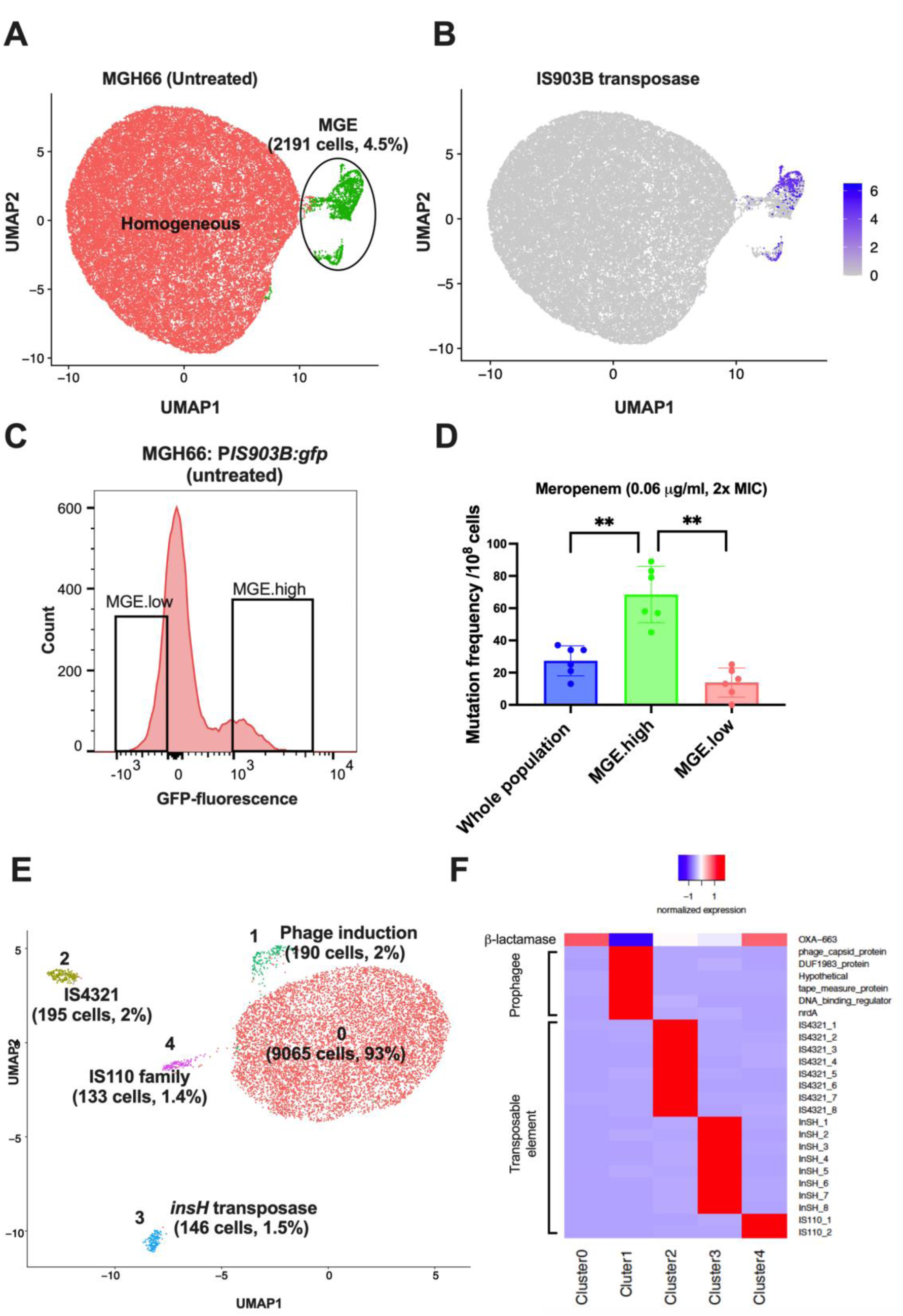
BacDrop reveals *within* population heterogeneity driven largely by mobile genetic elements (MGE). (A and B) In an untreated culture of MGH66, a population showing high-level expression of IS903B transposase (MGE, 4.5%; green) was detected. (C and D) Flow cytometry of reporter MGH66 strains expressing GFP driven by the promoter of *IS903B* (MGH66:P*IS903B:gfp*) shows a heterogenous expression pattern. The MGE.high population (∼10% of the whole population) and MGE.low population (∼10% of the whole population) were sorted into MHB medium without antibiotics, and mutation frequencies (D) were measured under meropenem treatment at the concentration of 0.06 μg/mL (2x MIC) using modified fluctuation analysis. Mutation frequencies of the MGE.high subpopulation were significantly higher (p= 0.002) than these of the MGE.low subpopulation and these of the unsorted controls (blue, whole population) All experiments in panel (C-D) were repeated three times with two biological replicates. Error bars were plotted as the standard deviation. The Student’s t-test was used for statistical analysis. (E and F) MGE-driven subpopulations were detected in another *K. pneumoniae* clinical isolate BIDMC35. UMAP (C) and heatmap (D) shows 4 subpopulations differing from the majority population (Cluster 0; red) of BIDMC35. Clusters 2, 3, and 4 are each driven by the high expression of a different transposase gene. In cluster 1, nearly all highly expressed genes belong to a prophage in BIDMC35 genome. Expression levels in all genes are normalized to expression in Cluster 0 (D).

Previously we had functionally shown that MGH66 and several other *K. pneumoniae* isolates have high-level transposon insertional mutagenesis activity, which contributes to their high frequencies of carbapenem resistance acquisition [38]; however, bulk RNA-sequencing had failed to show elevated expression of any transposon genes in these strains. Here, the existence of this subpopulation with high-level transposon expression provides a possible explanation for the strain’s elevated carbapenem-resistance frequencies, with resistance likely emerging from this small subpopulation. To test this hypothesis, we engineered a MGH66 reporter strain expressing the short-lived green fluorescent protein [39] driven by the promoter of one copy of the *IS903B* transpose genes (MGH66: P*IS903B:gfp*). Consistent with the BacDrop results, we observed heterogeneous expression pattern of *gfp* in this reporter strain by flow cytometry (Fig. 4C). We then performed fluorescence activated cell sorting (FACS) and sorted the MGE.high (high GFP expression) and MGE.low (low GFP expression) populations (Fig. 4C-4D) and measured their mutation frequencies under meropenem treatment using a modified fluctuation analysis [38]. The MGE.high population had at least 7 times higher mutation frequencies then the MEG.low population under meropenem treatment (Fig. 4D), confirming our hypothesis that resistance is more likely to emerge from this subpopulation highly expressing MGE genes and revealing phenotypic consequences of the different subpopulations.

To examine the robustness of the MGE subpopulation in our datasets and to compare the relative values of deeper sequencing of fewer cells versus more shallow sequencing of more cells, we analyzed the untreated samples in Replicate 1 and Replicate 2 libraries separately, along with a smaller BacDrop library containing ∼3,000 cells of MGH66 similarly collected but sequenced more deeply. We sequenced this smaller library to obtain 80,000 reads per cell with recovery of ∼2,000 cells with at least 15 mRNA genes per cell, and collectively an average of 85 mRNA genes detected per cell. Analysis of this smaller library detected the same MGE population identified in the larger cell population, albeit less distinctly (Fig. S4). In contrast, both the replicates of the larger libraries, even when sequenced only to a depth to obtain ∼30 mRNA genes per cell, identified an additional small subpopulation (0.25% - 0.36%) featuring high-level expression of maltose transport genes that was not identified in the smaller library, despite its deeper sequencing and higher coverage (Fig. S4). Together, these results show that analyzing larger numbers of cells can help to reveal heterogeneity within a bacterial population, which is consistent with analyses in eukaryotic systems [7, 19, 20, 25, 26, 28, 29]. Although the coverage can be higher when fewer cells were analyzed, increasing the scale, rather than the coverage, resulted in identification of a rare population.

To determine if MGE-driven subpopulations are unique to MGH66, we applied BacDrop to another *K. pneumoniae* clinical isolate (BIDMC35) (Table S2). BIDMC35 is a carbapenem-resistant isolate in which carbapenem resistance results from a transposon disruption of the major porin gene *ompK36* and the transposon-mediated high-level expression of a ß-lactamase gene *bla*_OXA-663_ [40]. We again observed MGE-driven subpopulations, as in MGH66. Analyzing 9,748 BIDMC35 single cells that passed the analysis threshold at a sequencing depth of ∼2,000 reads per cell (Fig. 4E-4F), we identified three clusters each driven by a unique transposon gene, including Cluster 2 driven by an IS4321 family transposase (195 cells, 2%), Cluster 3 driven by the *insH* transposase (146 cells, 1.5%), and Cluster 4 driven by an IS110 family transposase (133 cells, 1.4%). Together with the observation in MGH66, it reinforces the finding that variable expression of MGEs may be one of the major drivers of population heterogeneity.

In BIDMC35, besides these MGE-driven subpopulations, we observed another unique subpopulation (190 cells, 2%) driven by the 30- to 320-fold higher expression of a group of prophage genes (Fig. 4E-4F, Table S6), compared to the rest of the populations, indicating that this cluster of cells was likely undergoing spontaneous phage induction. This observation is similar to the phage-induction subpopulation reported in the microSPLiT study [16]. Additionally, we found the expression of *bla*_OXA-663_ is significantly lower in this phage-induction subpopulation (Fig. 4E), possibly caused by spontaneous phage induction.

### BacDrop reveals heterogeneous stress responses to antibiotic exposure

Finally, to determine if a perturbed population might have heterogenous dynamic transcriptional responses, we analyzed the single cell responses after different antibiotic exposures. Under the conditions in which these samples were generated, the numbers of cells queried, and the level of resolution they were analyzed, we did not detect obvious heterogeneity of response to ciprofloxacin or gentamicin. This contrasts with what we observed for meropenem treatment.

We had been particularly puzzled by the fact that despite having the same impact on cell killing as ciprofloxacin and gentamicin at 30 minutes (Fig. S5A), meropenem treatment only induced minimal transcriptional responses based on bulk RNA-seq analysis, in contrast to the other two antibiotics (Fig. 3B). We thus analyzed the transcriptional response at the single cell level for the meropenem-treated samples and indeed, in addition to the MGE-driven subpopulation, we found four interesting subpopulations with distinct molecular responses after meropenem treatment (Fig. 5A-5B; Fig. S5B). These subpopulations each showed co-upregulation of genes with four different, major biological functions, including: i) a stress response cluster featuring high-level expression of stress response genes such as *rseB, yidC,* and *yhcN*; ii) a cell wall/membrane synthesis cluster featuring highly expressed genes involved in cell wall and membrane synthesis, i.e., *nlpl* and *lpxH*; iii) a cell wall synthesis and DNA replication cluster featuring highly expressed genes involved in DNA synthesis and cell wall/membrane synthesis such as *dnaG* and *ftsI*; and iv) a cluster characterized by the upregulation of genes involved in cold shock response, such as *pnp* and *cspD*. Of note, although initially identified as a cold shock protein, CspD was reported to be a DNA-binding toxin that inhibits DNA synthesis and induces the formation of persisters in *E. coli* [41].

**Figure 5.**
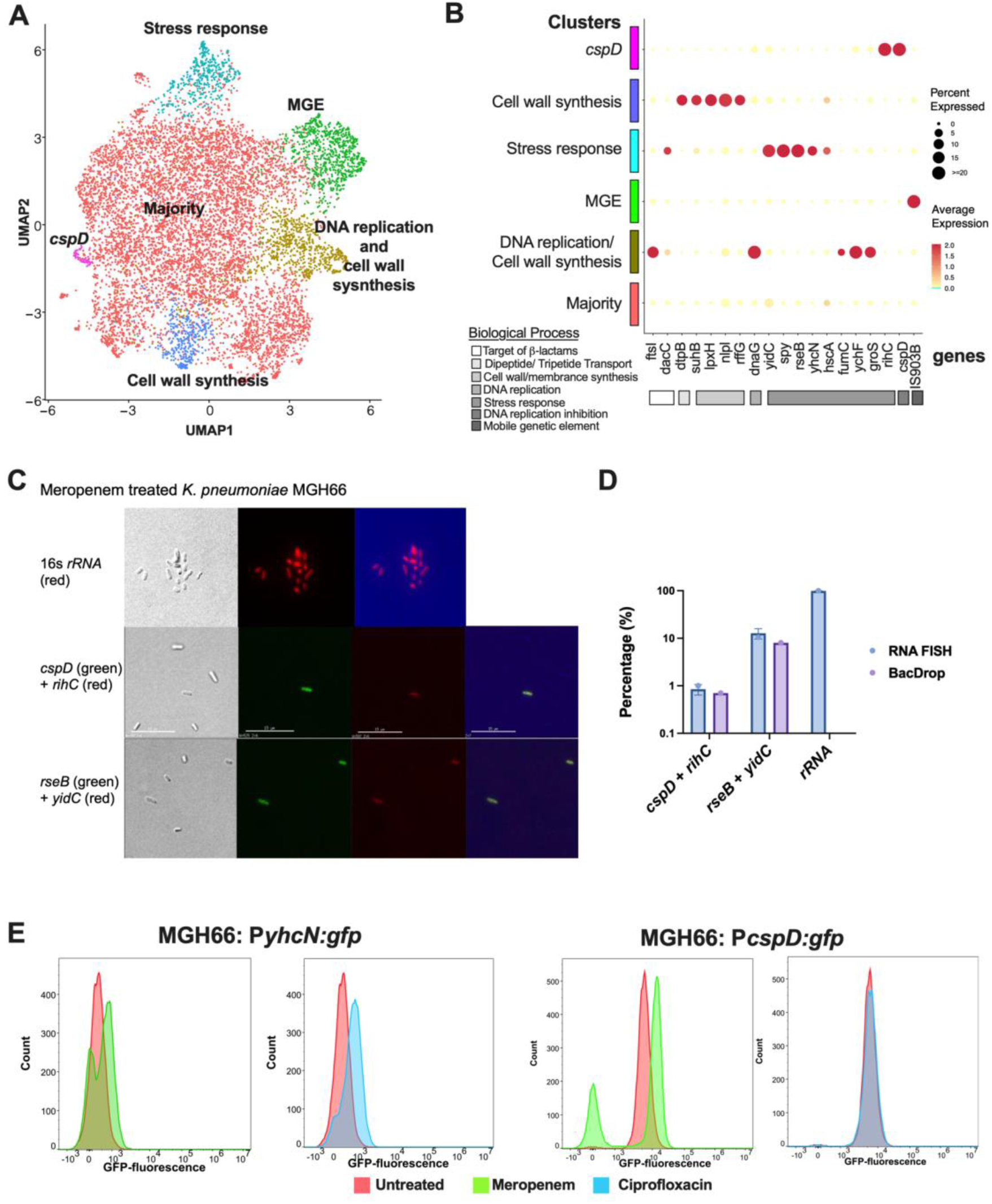
BacDrop reveals heterogeneous responses to meropenem exposure. (A) Besides the subpopulation highly expressing the IS903B transposase gene (MGE), meropenem treatment induced heterogenous responses, including a stress-response subpopulation, a cell wall synthesis subpopulation, a DNA replication and cell wall synthesis subpopulation, and a *cspD*-expressing subpopulation. (B) Dot plot showing the expression of genes that are significantly different among clusters and the percentage of cells expressing these genes in each cluster. (C) Validation of subpopulations identified in the meropenem- treated MGH66 using RNA FISH with double marker genes from the “*cspD* expressing” subpopulation (*cspD* + *rihC*) and the “strong stress response” subpopulation (*rseB* + *yidC*). A 16s *rRNA* probe was used as the positive control to show that more than 99% cells were successfully permeabilizated and hybridized. Subpopulations co-expressing double marker genes were identified. (D) Across 20 fields of view, the RNA FISH results showed that ∼0.85 % cells co-expressed *cspD* and *rihC*, and ∼12% cells co-expressed *rseB* + *yidC*. This result verified the BacDrop result in which ∼0.7% cells co-expressed *cspD* and *rihC,* and ∼8% cells co- expressed *rseB* + *yidC*. (E) Flow cytometry of reporter MGH66 strains expressing GFP driven by the promoter of *yhcN* (MGH66:P*yhcN:gfp*; left) or *cspD* (MGH66:P*cspD:gfp*; right) shows a heterogenous response to meropenem (green) but not to ciprofloxacin (blue) treatment, relative to untreated control (red). All experiments in panel (C-E) were repeated three times. Error bars were plotted as the standard deviation. The Student’s t-test was used for statistical analysis.

Having identified genes highly expressed in specific subpopulations after meropenem treatment (Fig. 5B, Table 1), we used RNA fluorescence *in situ* hybridization (FISH) to validate the finding of the “stress response” and “*csp*D-expressing” subpopulations. We simultaneously probed for two genes highly co-expressed in the “stress response” cluster, *rseB* and *yidC,* and confirmed the existence of this cluster (Fig. 5C). Similarly, we simultaneously probed for *cspD* and *rihC,* two genes highly co-expressed in the “*cspD-*expressing” cluster to confirm their existence (Fig. 5C). The percentage of cells making up each cluster as detected by RNA-FISH and BacDrop were very similar (Fig. 5D).

**Table 1.** Marker genes identified in clusters of the meropenem-treated sample.

| **Gene name** | **Gene product** | **Function** | **Biological pathways** | **Cluster** | **Log2 fold change** | **P value** |
| --- | --- | --- | --- | --- | --- | --- |
| *ftsI* | PBP3 | Target of β-lactams | Cell wall synthesis | DNA replication and cell wall synthesis | 5.2957 | <1.00E-300 |
| *dacC* | PBP6 | Target of β-lactams | Cell wall synthesis | Stress response | 3.7036599 | 1.81E-40 |
| *dtpB* | DtpB | Transportation of beta-lactams | Dipeptide/Tripeptide Transport | Cell wall synthesis | 6.281594 | <1.00E-300 |
| *suhB* | SuhB | Membrane protein transport and insertion | Cell wall/membrane synthesis | Cell wall synthesis | 4.406598 | 1.39E-125 |
| *lpxH* | LpxH | Lipid A biosynthesis | Cell wall/membrane synthesis | Cell wall synthesis | 7.605374 | <1.00E-300 |
| *nlpI* | NlpI | Cell wall synthesis, bind to PBP4 and PBP1A | Cell wall/membrane synthesis | Cell wall synthesis | 3.487722 | 2.54E-87 |
| *rffG* | RffG | LPS synthesis | Cell wall/membrane synthesis | Cell wall synthesis | 4.280285 | 5.97E-107 |
| *dnaG* | DnaG | DNA replication | DNA replication | DNA replication and cell wall synthesis | 4.490715 | <1.00E-300 |
| *yidC* | YidC | Membrane damage response | Stress response | Stress response | 4.0687447 | 8.25E-100 |
| *spy* | Spy | Membrane damage response | Stress response | Stress response | 4.0142941 | 2.33E-98 |
| *rseB* | RseB | Membrane damage response | Stress response | Stress response | 5.1874707 | 2.26E-218 |
| *yhcN* | YhcN | Cytoplasmic acid stress | Stress response | Stress response | 6.1654092 | <1.00E-300 |
| *hscA* | HscA | Cold shock | Stress response | Stress response | 3.2262902 | 5.75E-26 |
| *fumC* | FumC | Oxidative stress | Stress response | DNA replication and cell wall synthesis | 5.304257 | <1.00E-300 |
| *ychF* | YchF | Oxidative stress | Stress response | DNA replication and cell wall synthesis | 5.286679 | <1.00E-300 |
| *groS* | GroES | Stress response | Stress response | DNA replication and cell wall synthesis | 4.510794 | 2.41E-261 |
| *rihC* | RihC | DNA damage response | Stress response | *cspD* expressing | 6.16122 | <1.00E-300 |
| *cspD* | CspD | Persister formation | DNA replication inhibition | *cspD* expressing | 6.342385 | <1.00E-300 |
| *IS903B* | IS903B | Insertion sequence | Mobile genetic elements | MGE | 4.463117 | <1.00E-300 |

We also used fluorescence cytometry to validate the expression patterns of genes in the “stress response” and “*cspD-*expressing” clusters. We engineered MGH66 reporter strains expressing the short-lived green fluorescent protein[39] driven by the promoter of *yhcN,* a “stress response” gene (MGH66:P*yhcN:gfp*), or *cspD* (MGH66:P*cspD:gfp*). Indeed, consistent with BacDrop results, meropenem but not ciprofloxacin induced heterogeneous expression of both *yhcN* and *cspD* (Fig. 5E). We confirmed that the cells with reduced fluorescence of MGH66:P*cspD:gfp* (GFP-low) were live and not simply dying using a live-dead stain and plating for colony forming units (Fig. S6). Interestingly, the GFP-low population actually had lower fluorescence than the untreated population, suggesting that in fact, there may be suppression of *cspD* expression in this population. Of note, the fractions of cells highly expressing these two genes were greater by flow cytometry than by BacDrop; this would be consistent with the fact that RNA and protein levels are not necessarily well correlated in single cells [13], as one copy of mRNA can produce one to hundreds of copies of proteins and mRNA is less stable than protein. Nevertheless, the heterogeneous responses were observed by both methods of measurement.

Taken together, under meropenem perturbation, we observed strong heterogeneous responses driven by various stress response pathways that had been previously masked in bulk RNA-seq results. Rather than uniformly turning on a specific stress response pathway in all cells, a diverse range of stress responses appear to be induced in different subpopulations, which could potentially contribute to heterogeneous cell fates such as cell lysis or antibiotic tolerance.

### BacDrop identifies a subpopulation with reduced meropenem efficacy and increased persisters

Given CspD’s reported role in inducing the formation of antibiotic-tolerant persisters in *E. coli* [41], we wondered whether antibiotic tolerant cells might make up a greater percentage of the GFP-high than the GFP-low subpopulation, with cells in which *cspD* had been induced surviving preferentially under meropenem treatment. Because CspD plays a role in persister formation rather than maintenance, we anticipated that some cells might induce *cspD* and thus make high levels of GFP, become antibiotic tolerant, then turn off CspD expression resulting in reduced expression of GFP as the GFP is degraded. Despite this possibility, we nevertheless hypothesized that the GFP-high subpopulation might still be enriched for tolerant cells compared to the GFP-low subpopulation. We performed FACS with the MGH66 GFP transcriptional reporter strain (MGH66:P*cspD:gfp*) coupled with dead-cell stain, after a 30-minute exposure to meropenem. We sorted live MGH66:P*cspD:gfp* cells into the two subpopulations, GFP-high and GFP-low (Fig. 6A), directly into liquid media with and without meropenem. (We confirmed that all sorted cells have similar survival rates on LB agar plates without antibiotics; Fig. S5.) We then plated an aliquot of the cells from the resulting 4 samples onto solid agar without antibiotics to enumerate surviving bacteria over time. Indeed, the GFP-high subpopulation was enriched for meropenem tolerant cells compared to the GFP-low subpopulation, with evidence of a persister population in the GFP-high but not the GFP-low subpopulations (Fig. 5B). We verified that no genetic mutations were acquired by these persister cells via whole genome sequencing. Consistent with this observation, we also observed ∼ 100 times more persister cells in a MGH66 strain over-expressing *cspD*, though the over-expression of *cspD* did not reduce antibiotic susceptibility as reflected in the minimal inhibitor concentration of meropenem (Fig. 5C-5D). Together, these results confirmed the role that *cspD* plays in persister formation in a subpopulation of *K. penumoniae* MGH66.

**Figure 6.**
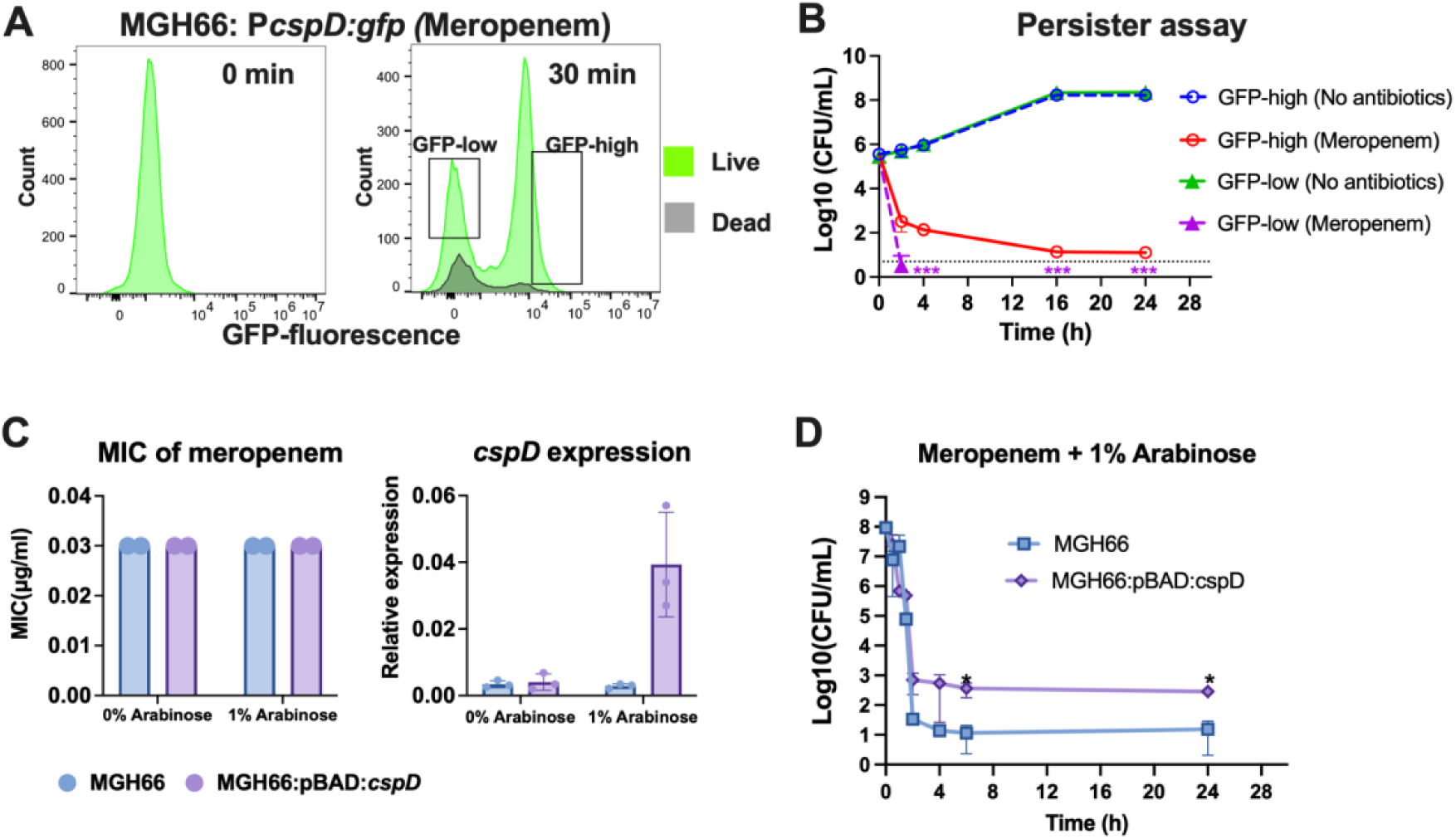
*cspD*-expressing cluster was enriched with persister cells. (A) MGH66:P*cspD:gfp* cells were analyzed by FACS at time 0 and 30 minutes after exposure to meropenem (2µg/mL). After 30 minutes, live cells from GFP-low and GFP-high subpopulations were identified and sorted. (B) GFP-low and GFP-high subpopulations sorted from meropenem- treated MGH66:P*cspD:gfp* cells differed in their response to meropenem. GFP-low and GFP- high subpopulations were sorted directly into media with no antibiotic or with meropenem (2 µg/mL). Samples were then taken over time and plated on solid agar to enumerate CFU. Persisters only emerged from the GFP-high subpopulation. The limit of detection is indicated by black dashed line. Three asterisks from the treated GFP-low subpopulation indicate either no persister was present or the numbers of persisters were below the limit of detection. (C-D) Overexpression of *cspD* in MGH66 increased numbers of persisters but did not affect the susceptibility of meropenem. *cspD* driven by the arabinose inducible promoter pBAD was transformed into MGH66. (C) Induction of *cspD* with 1% arabinose did not affect the minimal inhibitory concentration (MIC) of meropenem. (D) When *cspD* was induced with 1% arabinose and treated with meropenem at 2 µg/mL (purple), a greater number of persister cells were formed compared to the culture without arabinose induction under 2 µg/mL meropenem treatment (blue). All experiments were repeated three times. Error bars were plotted as the standard deviation. The Student’s t-test was used for statistical analysis.

## Discussion

We report a novel bacterial scRNA-seq method, BacDrop, that is robust and reproducible, and leverages droplet-based technology to enable the massively parallel profiling of the transcriptional program of thousands to millions of single bacterial cells. We applied BacDrop to study the naturally occurring heterogeneity in a heretofore presumed uniform culture of bacteria and its heterogenous responses to perturbation. Due to limited scale, previous genome-wide single cell transcriptional studies on smaller numbers of cells (hundreds to thousands) have to date focused predominantly on samples with artificially created heterogeneity, created as mixes of different populations, for method demonstration of *between* population heterogeneity [14–16]. Here, we characterized both a stable and a dynamic population of cells derived from the same bacterial isolate and demonstrated a diversity of states *within* the stable population and heterogenous responses in the dynamic population after perturbation with antibiotic.

We have shown that BacDrop can be applied to a wide range of numbers of cells, from thousands to millions. Importantly, we demonstrated that studying large numbers of cells can more robustly identify population heterogeneity including rare subpopulations even with fewer mRNA genes per cell (Fig. S4). This result is in line with analyses in mammalian systems, suggesting that the ability to analyze a large number of cells is important for the identification of rare cell types and states, particularly in light of the relatively sparse per cell coverage afforded by single cell technologies [2, 7, 10, 19, 20, 25–29, 42]. With the ability to characterize large numbers (millions) of cells without requiring any prior knowledge of the genomes of interest, the limitation for understanding the single cell transcriptional programs within complex bacterial communities is no longer technical. Instead, one of the major limitations becomes that of sequencing cost, which currently necessitates a compromise between sequencing large numbers of cells and the sequencing depth of individual cells, with the field currently favoring large numbers of cells. With the rapid development of sequencing technologies and platforms, one can easily imagine that such compromises may no longer be required in the near future.

Using BacDrop, we report the observation of *within* population heterogeneity driven predominantly by the expression of MGEs in the presumed uniform cultures of *K. pneumoniae*. While MGEs drive genetic diversity by their relatively random movement in the genome, we find that the expression levels of individual MGEs are also highly variable within populations of MGH66 and BIDMC35 (Fig. 4). Whether this heterogeneity in expression is due to genotypic variation resulting from the movement of genetic elements such as transposon insertions or phase variation, to transcriptional changes in response to stochastic/or local microenvironmental cues, or to epigenetic mechanisms, remains to be understood. However, the consequences of such variation can be significant. We demonstrated that the high-level expression of MGE genes found only in a subpopulation of MGH66 contribute to the strain’s propensity to become carbapenem resistant, consistent with our previous finding that high-level transposon mutagenesis plays an important role in high-frequency evolution of carbapenem resistance [38].

Meanwhile, in response to a perturbation such as exposure to the important antibiotic meropenem, BacDrop revealed a wide range of transcriptional responses that are masked in bulk RNA-seq. Interestingly, a diverse range of stress responses appear to be induced in different subpopulations rather than a single or uniform response occurring in all cells; this subpopulation diversity could potentially contribute to heterogeneous cell fates or phenotypic outcomes, including cell lysis or antibiotic tolerance. It remains to be understood whether this heterogeneous stress response occurring in subpopulations is a general phenomenon in response to stressors or is specific to certain types of stresses such as meropenem, which may be working in a more pleiotropic manner than is assumed, thereby eliciting a wide range of stress responses. Meanwhile, we examined one such subpopulation defined by high-level expression of *cspD,* a gene encoding a toxin that inhibits DNA replication and induces the formation of persisters, but that is not significantly upregulated in bulk RNA-seq results. Using both FISH and flow cytometry, we confirmed that *cspD* is indeed induced in a subpopulation after meropenem treatment. Moreover, we found a higher survival rate under meropenem treatment in the *cspD*-induced subpopulation, pointing to its role in antibiotic tolerance within this subpopulation and more generally, to the importance of certain responses in some subpopulations in surviving the lethal effects of antibiotics.

We have developed BacDrop to be a versatile method to characterize the transcriptional programs of thousands to millions of bacterial cells in multiple species. To demonstrate its utility, we validated BacDrop in four different pathogenic bacterial species. We expect that BacDrop should be easily adapted to more species, including commensal bacterial strains of the microbiome and other environmental species. Therefore, BacDrop will enable the identification of both heterogeneity in cell types (species) in a mixed community or cell states in a single strain under stable or dynamic conditions. We thus propose that BacDrop will become a powerful tool for a broad range of studies, including studies focused on elucidating phenotypic heterogeneity, understanding bacterial interactions in microbial communities, dissecting host-pathogen interactions, expanding our knowledge of the microbiome beyond genomes and bulk metabolomics, and investigating the emergence of antibiotic resistance, persistence and tolerance.

## Acknowledgments

We thank Jonathan Livny, Noam Shoresh, and Eugenio Mattei for the valuable discussions on the data analysis, Zohar Bloom-Ackermann for the suggestions on the flow cytometry, Anne Clatworthy and Thulasi Warrier for the comments on the manuscript, Kevin Grosselin and Ming Pan for their discussion on the droplet generation steps. We thank Aviv Regev for her expert advice and comments on the manuscript. We thank the Microbial Omics Core at the Broad Institute for generating bulk RNA-seq libraries, and thank the Flow Cytometry Core Facility at the Broad Institute for cell sorting. This publication was supported in part by the National Institute of Allergy and Infectious Diseases of the National Institutes of Health under award 5R01AI117043-05 to DTH, and by a generous gift from Anita and Josh Bekenstein.

## Author contributions

P.M. and D.T.H. designed the study and wrote the manuscript. P.M. and L.L.H performed the experiments. P.M., H.M.A. and S.J.G. performed data analysis. R.P.B. provided extensive input and suggestions on the design of this study and manuscript preparation. R.N. provided extensive input and suggestions on the development of BacDrop. C.S. provided extensive input and suggestions on the data analysis, cluster identification and manuscript revision. All authors have read and approved the manuscript.

## Methods

### Bacterial strains and culture

Bacterial strains used in this study are listed in Table S2. *K. pneumoniae*, *E. coli*, and *P. aeruginosa* were cultured in Luria-Bertani (LB) medium or Mueller-Hinton Broth (MHB) with shaking at 37 °C. *E. faecium* was cultured in Todd-Hewitt broth (THB) with shaking at 37 °C. For the GFP experiment, *E. coli* strains expressing *gfp* driven by different promoters were cultured separately in LB at 37 °C. Early exponential growth phase cells (OD_600_ ∼0.2) were collected and fixed immediately. For the antibiotic treatment experiment, *K. pneumoniae* clinical isolate MGH66 was cultured in MHB at 37 °C until early exponential phase. Then the culture was diluted to OD_600_ 0.05 in MHB. After growing for two doublings (∼ 40 min, OD_600_ ∼0.2), the cultures were split into four equal volume cultures. One culture was left untreated, while the other three were treated with relevant antibiotic at breakpoint concentrations set by the Clinical and Laboratory Standards Institute (CLSI): 2 µg/mL for meropenem, 1 µg/mL for ciprofloxacin and 4 µg/mL for gentamicin. After 30 min, 7 mL of cells were collected from each of these cultures, and immediately proceeded with the cell fixation protocol. For the bulk RNA-seq experiments, samples were collected using the same treatment schemes, and three biological replicates were included in each condition. For the BIDMC35 experiment, BIDMC35 was cultured in LB medium at 37 °C until early exponential phase (OD_600_ ∼0.2). 7 mL cells were then collected and immediately proceed with the cell fixation protocol.

### Cell fixation and permeabilization

All reagents (Table S7) for cell fixation and permeabilization were kept ice-cold. Up to 10 billion bacterial cells grown at specified conditions were collected by centrifuging at 5525 x g for 10 min at 4 °C. The supernatant was removed and cell pellets were resuspended in 7 mL fresh, ice-cold 4% formaldehyde (Sigma, CN 47608) in 1x PBS and incubated with shaking overnight at 4 °C (Digital Platform Rocker Shaker, VWR). Following overnight fixation, cells were centrifuged at 5525 x g for 10 min at 4 °C. The supernatant was removed and cells were resuspended in 7 mL PBS-RI (1x PBS supplemented with 0.1 U/µL NxGen RNase inhibitor (Lucigen, CN 30281)). Cells were centrifuged again at 5525 x g for 10 min at 4 °C and pellets were resuspended in 700 µL PBS-RI. Subsequent centrifugations were carried out at 7000 x g for 5 minutes at 4 °C. We noticed that some species, e.g., *P. aeruginosa*, do not pellet very well at this centrifugation speed (Fig. S1F). Thus, we recommend optimizing the centrifugation speed for the specific species of interest. Cells were centrifuged and resuspended in 700 µL 50% Ethanol in PBS-RI. Cells were then washed twice with PBS-RI. After the second wash, cells were resuspended in 1 mL 100 mM Tris-HCL (pH 7.5) supplemented with 0.1 U/uL NxGen RNase inhibitor. Cells were diluted 100x and quantified using a hemocytometer (VWR, CN 102966).

We found that the number of cells in a permeabilization reaction is critical for achieving sufficient permeabilization. Insufficient permeabilization may result in inefficient rRNA depletion and gDNA removal. For 1 permeabilization reaction, up to 40 million cells were centrifuged and resuspended in 250 µL 0.04% Tween-20 in 1x PBS. If more cells are desired, multiple parallel reactions can be set up. Immediately following a 3-minute incubation on ice, 1 mL cold PBS-RI was added, and cells were spun down and resuspended in 200 µL lysozyme mix (100 mM Tris (pH 8.0), 50 mM EDTA pH 8.0, 0.25 U/µL NxGen RNase Inhibitor, 2.5 mg/mL lysozyme). Cells were then incubated at 37 °C for 15 minutes. After the incubation, 1 mL PBS-RI was added and cells were washed twice with 175 µL PBS-RI. After the second wash, cells were resuspended in 150 µL PBS (without RNase inhibitor added) and cell concentrations were measured.

### In-cell rRNA depletion and gDNA removal

Immediately after cell permeabilization, up to 40 million cells were centrifuged and resuspended in 11 µL nuclease-free H_2_O. The cell number is critical for achieving efficient rRNA depletion (Fig. S1E). 2 µL NEBNext Bacterial rRNA depletion solution and 2 µL Probe Hybridization Buffer (NEB, CN E7850) were mixed with cells on ice. The hybridization was conducted per the following (lid temperature set to 55 °C): 50 °C for 2 minutes, ramp down to 22 °C at 0.1 °C /second, and hold at 22 °C for 5 minutes. Probe hybridization was immediately followed by RNase H digestion by mixing the probe-hybridized cells with 2 µL RNase H reaction buffer, 2 µL Thermostable RNase H (NEB, CN E7850), and 1 µL nuclease-free H_2_O, followed by a 30-minute incubation at 50 °C (lid temperature set to 55 °C). The 20 µL reaction was centrifuged and resuspended in 10 µL DNase-RI buffer (1 µL DNase I reaction buffer (Sigma, CN AMPD1), 1 µL DNase I (Sigma, CN AMPD1), 0.025 µL NxGen RNase inhibitor (Lucigen, CN 30281), 8 µL nuclease-free H_2_O). The reaction was incubated at room temperature for 30 minutes, and the DNase treatment was stopped by adding 1 µL Stop Solution (50 mM EDTA) and incubating at 50 °C for 10 minutes. After the incubation, cells were centrifuged and washed twice with 100 µL PBS-RI. After the second wash, cells were resuspended in 20 µL 0.5x PBS supplemented with 1 U/µL SUPERase-In RNase inhibitor and used immediately for in-cell reverse transcription.

### In-cell reverse transcription, round 1 cell barcoding and sample multiplexing

The round 1 plate barcoding and sample multiplexing is achieved via RT reactions in 384- or 96- well plates. 384 RT primers (Table S1) containing UMI sequences and round 1 plate barcodes (CB1) were synthesized at Integrated DNA Technologies at 100 µM concentration. The primers were diluted with ddH_2_O to a working concentration of 25 µM, and 2.5 µL of each primer was aliquoted into individual wells of the 384- or 96- well plates. The rRNA and gDNA depleted cells was diluted with nuclease-free H_2_O supplemented with 1 U/µL SUPERase-In RNase inhibitor and added to the plate containing RT primers (1 µL cells per well). Then the cell-primer mix was incubated at 55 °C for 5 minute and immediately put on ice. For each well, the RT master mix, containing 0.25 µL DTT (100 mM), 0.25 µL dNTP (10 mM each), 0.25 µL SUPERase-In, 1 µL RT buffer, 0.25 µL Maxima H Minus reverse transcriptase (Thermo Scientific, CN EP0753), was added and the RT reaction was incubated as follows (set lid temperature to 60 °C): 22 °C for 30 min, 50 °C for 10 min, 3 cycles of [8 °C for 12 s, 15 °C for 45 s, 20 °C for 45 s, 30 °C for 30 s, 42 °C for 2 min, 50 °C for 3 min], 50 °C for 5 min, hold at 4 °C.

### In-cell cDNA 3’ poly-A tailing

After RT, cells were recovered and pooled from 384- or 96- well plate. For multiplexed samples, we suggest to only pool cells from the same sample together, generating individual pools for each sample. This will allow more flexibility to adjust cell numbers of different samples during droplet generation. Then the pooled cells were centrifuged and each pool was resuspended 40 µL nuclease-free H_2_O supplemented with 1 U/µL SUPERase-In. The poly-A tailing reaction was set up as the following: 38 µL cells, 5 µL 10x terminal transferase buffer, 1 µL dATP (100 mM) (NEB, N0446S), 1 µL SUPERase-In, 5 µL terminal transferase (TdT) (NEB, CN M0315L). The reaction was incubated at 37 °C for 1 hour. Then 10 µL 0.2 M EDTA was added to each 50 µL reaction and incubated at room temperature for 10 minutes. Cells were then centrifuged and resuspended in 10 µL nuclease-free H_2_O supplemented with 1 U/µL SUPERase-In. Cells were diluted and concentrations were quantified.

### Droplet generation

The Chromium Next GEM Chip H (10x Genomics, PN 1000161) and Chromium Next GEM Single Cell ATAC Library & Gel Bead Kit (10x Genomics, PN 1000176) was used for the droplet generation. The unused wells were filled with 70 µL (row 1), 50 µL (row 2), and 40 µL (row 3) 50% glycerol solution. The desired number of cells was diluted to 33.75 µL with nuclease-free H_2_O supplemented with 1 U/µL SUPERase-In. Right before loading the chip, 33.75 µL cells were mixed with a PCR master mix containing 37.5 µL KAPA HiFi HotStart ReadyMix (Roche, CN KK2602), 2.25 µL second strand synthesis primer SMRT_dT (10 µM) (Table S8), and 1.5 µL Reducing Agent B (10x Genomics, PN 2000087). The chip was loaded with 70 µL PCR-cell mix (row 1), 50 µL Gel Beads (row 2), and 40 µL partitioning oil (row 3), and run on the Chromium system. Approximately 100 µL droplet emulsion was obtained in row 3.

### Second strand cDNA synthesis and round 2 droplet barcoding in droplets

To increase thermostability of the droplets, we split each 100-µL emulsion into 4 25-µL reactions into PCR tubes (USA Scientific, CN 1402-4700). In each reaction, 25 µL 5% FC-40 oil (RAN Biotechnologies, CN 008-FluoroSurfactant-5wtF) was added to the bottom and 50 µL mineral oil (Sigma, CN M5904) was added on the top. The 2^nd^ strand cDNA synthesis and round 2 cell barcoding was performed as follows: 95 °C for 30 s, 39 °C for 5 min, 65 °C for 10 min; then 4 cycles of [98 °C for 20 s, 62 °C for 15 s, 72 °C for min], 72 °C for 5 min, hold at 4 °C.

### Breaking emulsions and cDNA purification

After the round 2 cell barcoding was finished, the mineral oil and FC-40 oil was removed from the PCR tube, being careful and not to remove the middle layer which contains the emulsion. The emulsion was then combined. In cases where only a small number of cells are desired for library construction, the emulsion can be kept separately but all downstream reaction volume should be reduced accordingly. Each 100 µL emulsion can then be broken by adding 125 µL Recovery Agent (10x Genomics, PN 2000087). The tubes were inverted 10 times and centrifuged briefly to ensure that all droplets were broken. 125 µL Recovery Agent/Partitioning Oil (pink) from the bottom of the tube was then removed. cDNA was first purified using Dynabeads. In brief, for each reaction, a mix containing 182 µL cleanup buffer (10x Genomics, PN 2000088), 8 µL Dynabeads MyOne SILANE (10x Genomics, PN 2000048), 5 µL Reducing Agent B, and 5 µL Nuclease-free water was added. After mixing and incubating at room temperature for 10 minutes, samples were placed on a magnetic separator and washed twice with freshly prepared 80% ethanol. After removing ethanol from the second wash, each sample was eluted in 40.5 µL elution buffer that was prepared by mixing 98 µL Buffer EB (Qiagen, CN 19086), 1 µL 10% Tween 20 (Teknova, CN T0710), and 1 µL Reducing Agent B. Then 40 µL of the elution was transferred to a fresh PCR tube and subjected to a 0.6x Cleanup with AMPure XP beads (Beckman Coulter, CN A63881). The cDNA was then eluted in 30 µL nuclease-free water.

### cDNA enrichment

Before the cDNA enrichment, a qPCR reaction, containing 1 µL cDNA, 5 µL KAPA HiFi HotStart Ready Mix, 0.3 µL primer P5 (10 µM), 0.3 µL primer SMRT_PCR (10 µM) (Table S8), 2 µL 5x SYBR green (VWR, CN 12001-796), and 1.4 µL nuclease-free water, was set up to determine cycle numbers of the cDNA enrichment in a real-time thermocycler using the following program: 98 °C for 3 min, 30 cycles of [98 °C for 20 s, 67 °C for 20 s, 72 °C for 3 min], 72 °C for 5 min, hold at 4 °C. The cycle numbers at which the qPCR reaction reaches early exponential amplification phase was determined as the cycle numbers for cDNA enrichment. For cDNA enrichment, 25 µL of cDNA was mixed with 125 µL KAPA HiFi HotStart Ready Mix, 7.5 µL primer P5 (10 µM), 7.5 µL primer SMRT_PCR (10 µM), and 85 µL nuclease-free water. The cDNA was enriched using the same program as the qPCR reaction, using the cycle numbers determined from the qPCR reaction. The enriched cDNA was purified using 0.6x AMPure XP beads and eluted in 50 µL nuclease-free water.

### Illumina sequencing library construction

The Illumina Nextera XT DNA library preparation kit (Illumina, CN FC-131-1096) was used to prepare sequencing libraries with these following modifications: ∼ 2 ng enriched cDNA and 4 µL ATM was used for each 50 µL tagmentation reaction. Following tagmentation, 2.5 µL primer P5 (5 µM) and 2.5 µL Index 1 primer (N7**, Illumina, CN FC-131-2001) was used for the PCR enrichment. The PCR reaction was removed from the thermocycler and immediately put on ice after 5 cycles of PCR amplification. Then a qPCR reaction, containing 5 µL Nextera XT PCR reaction, 3 µL NPM, 1 µL primer P5 (5 µM), 1 µL Index 1 primer, 3 µL SYBR green, and 2 µL nuclease-free water, was set up to determine the remaining cycle numbers of the library enrichment using the following program: 72 °C for 3 min, 95 °C for 30s, 25 x [95 °C for 10 s, 55 °C for 30 s, 72 °C for 30 s], 72 °C for 5 min, hold at 10 °C. The cycle numbers at which the qPCR reaction reaches one third of the fluorescence saturation was determined as the remaining cycle numbers for library enrichment. Then the remaining 45 µL library enrichment PCR reaction was put back to a thermocycler and amplified using the same program with cycle numbers determined from the qPCR. The libraries were then subjected to a 0.6x cleanup with AMPure XP beads and eluted in 25 µL nuclease-free water.

### Illumina sequencing of BacDrop libraries

Libraries were diluted to desired concentrations and sequenced on the Illumina NovaSeq 6000 platform with standard sequencing primers, using the following specifications: Read 1: 60 bp; Read 2: 39 bp; Index 1: 8 bp; Index 2: 16 bp. Depending the scale of the experiment and the sequencing depth desired, NovaSeq 6000 SP (Illumina #20027464), S1 (Illumina #20012865), or S2 (Illumina #20012862) reagent kits were used.

### Bulk library construction

To construct sequencing libraries from bacterial cultures, cell pellets collected from 1 mL early exponential phase cultures were re-suspended in 500 µL TRIzol Reagents (ThermoFisher, CN 15596026) and frozen at -80 C for at least 20 min. Cells were then thawed and mixed with 250 µL of 0.1 mm diameter Zirconia/Silica beads (BioSpec Products), and lysed mechanically via bead-beating for 90 second at 10 m/sec on a FastPrep (MP Bio). After addition of 0.1 mL chloroform, each sample tube was mixed thoroughly by inversion, incubated for 3 minutes at room temperature, and centrifuged at 12,000 xg for 15 minutes at 4°C. The aqueous phase was mixed with an equal volume of 100% ethanol, transferred to a Direct-zol spin column (Zymo Research, CN R2051), and RNA was extracted according the Direct-zol protocol. The sequencing libraries were then generated using the RNAtag-Seq protocol[43].

To construct sequencing libraries from fixed cells, or fixed and permeabilized cells, 20 µL cells were pelleted and resuspended 20 µL lysis buffer (50 mM Tris pH 8.0, 200 mM NaCl, 25 mM EDTA pH 8.0) supplemented with 1.6 µL proteinase K (50 mg/mL). Cells were lysed at 55 °C for 1 hour, then RNA was purified using 1.5x AMPure RNAClean XP beads (Beckman Coulter, CN A63987) and eluted in 20 µL nuclease-free water. The sequencing libraries were then generated using the RNAtag-Seq protocol.

### Estimation of mRNA copy numbers in *E. coli* GFP strains using RT-qPCR

The estimation of mRNA copy numbers was performed using a protocol modified from a previous study[13]. In brief, a *gfp* gBlock fragment was synthesized at Integrated DNA technology. A serial dilution was performed to create *gfp* dsDNA standards ranging from 1 to 10^10^ molecules/µL. One µL of the standard was used in a 10 µL qPCR reaction to generate the standard curve. 10 ng RNA from each GFP strain (∼66,666 cells with the assumption that there is 0.15 pg RNA per bacterial cell) was converted into cDNA and subjected to qPCR reaction together with the *gfp* dsDNA standards. GFP mRNA copy numbers were estimated by mapping to the standard curve.

### Construction of GFP reporter strains

We amplified the promoter region of *cspD* or *yhcN* from MGH66 and ligated it into pUA139[39] using the BamHI and XhoI sites, resulting in pUA139-PcspD and pUA139-PyhcN. For the promoter of IS903B, we synthesized the promoter sequence at IDT with BamHI and XhoI restriction digestion sites as the following sequences and ligated it into pUA139: GGATCCAGAAATTCTCTGTTCCATGGTAGATTAATAAGTCCCCAACATTTAAATATACAGGATAATCTAAATATTACTTCGTTCTTATCCTTAATAAATGGCAAAATTTCATTTAATTTATTTTTCAAATTATTCTGATGCATGAGTTACCCTATAATTTACACATAAAGAAGGCTTTGTTGAATAAATCGAACTTTTGCTGAGTTGCTCGAG.

The construct was then transformed into *E. coli* DH5α and the plasmids were extracted and verified using Sanger sequencing. Electrocompetent cells of MGH66 was made as previously reported [38]. The extracted plasmids were then transformed into MGH66 via electroporation, generating MGH66: P*IS903B:gfp*, MGH66: P*cspD:gfp* and MGH66: P*yhcN:gfp*.

### Cell sorting of untreated MGH66: P*IS903B:gfp* and measurement of mutation frequencies

For cell sorting of MGH66: P*IS903B:gfp*, the top 10% cells with high-level GFP expression and bottom 10% cells with low-level GFP expression were sorted into MHB medium without any antibiotics. The sorted cells were then used as the starting culture for measuring the mutation frequencies. We used a modified fluctuation analysis to measure mutation frequencies as previously described [38]. In brief, ∼100 sorted cells were seeded into each well of 384-well plates, followed by incubating at 37C for 3 hours. After the incubation, cells from three randomly chosen wells were taken and plated for cell counting. The rest of the wells were added meropenem at 0.06 ug/mL (2x MIC) and treated overnight at 37C in a humidified chamber. The 2^nd^ day morning, wells with resistant mutants growing up were counted to calculate the mutation frequencies. An unsorted culture was included as a control. This experiment was repeated 3 times with 2 biological replicates each time.

### Flow cytometry of the meropenem-treated samples

The cells for flow cytometry were prepared using the same scheme as the meropenem-treated sample for BacDrop. After treating with meropenem at 2 µg/mL for 30 minutes, cells were diluted into PBS at the final concentration of 10^6^ cells/mL and immediately run through flow cytometer. For the flow cytometry including dead-cell staining, 10 µL propidium iodide (1 mg/mL) (Invitrogen, cat# P1304MP) was mixed with 1 mL PBS. Cells after treatment were then diluted in the staining buffer at the final concentration of 10^6^ cells/mL, and incubated in the dark at room temperature for 15 minutes followed by running through flow cytometer. To sort GFP-low and GFP-high populations, cells were run through fluorescence-activated cell sorting (FACS). We sorted ∼10^6^ cells into 2 mL LB without or with meropenem (2 µg/mL) from each population. A 50 µL aliquot from cells sorted in LB medium without antibiotics was immediately diluted and plated on LB agar plates without antibiotics. The colony forming unites (CFU) on LB agar plates were enumerated to calculate the survival rates. The rest of the cells was used to measure MICs using the standard microdilution protocol[44] or proceeded with the persister assay. For the persister assay, cells were cultured at 37 °C with shaking. At each time points, a 50 µL aliquot was taken, diluted and plated on LB agar plates without antibiotics. Three replicates were performed in each experiment.

### RNA FISH

RNA FISH probes were designed using Design Probes tool of DECIPHER[45]. 3 - 5 probes were designed for each target gene (Table S9). Probes were synthesized at Integrated DNA technologies and labeled with either Cy3 or Alex488 dye at the 5’. Probes of the same gene were pooled together and diluted into 10 µM stocks. Meropenem-treated MGH66 cells were fixed and permeabilized using the same protocol as the cell fixation and permeabilization steps in BacDrop. The hybridization was carried out in 40% hybridization buffer at 50 °C overnight and the following washing steps of washing were performed as described previously[46]. Cells were imaged using the DeltaVision widefield deconvolution imaging system with a 60x objective. ImageJ was used for image data analysis and cell quantification.

### Computational methods Processing of sequencing data

To build input matrices with gene-barcode information, we created a pipeline that pulls UMI, barcodes 1 and 2. In experiments where multiple samples were pooled for droplet barcoding, a demultiplexing step based on CB1 was performed to parse different samples. Threshold of valid cell barcodes, removal invalid cell barcodes, and final count table generation was performed using UMI-tools[47] (10.1101/gr.209601.116). Alignments were performed using BWA[48] and annotation of bam files were done using FeatureCounts[49].

### Analysis of BacDrop data

Once the count tables were made, we used the standard workflow of the R package Seurat 3[36] (https://doi.org/10.1038/nbt.4096, https://doi.org/10.1016/j.cell.2019.05.031) (v.3.2.2). We excluded genes that were not expressed in any cells in the dataset, and excluded cells that had fewer than 10 or 15 genes detected and cells that had abnormally high numbers of mRNA detected. For experiments done using MGH66, we identified three genes that showed consistently high-level expression in the majority of cells and in various conditions: WP-004174069.1, WP- 004174069.1-2, and WP-002920103.1. We use these three genes as internal controls and removed cells with a normalized expression of any of these three genes that is less than 50. The standard Seurat workflow prior to clustering was used including global normalization, feature selection, and scaling of gene expression. We used the top 2000 highly variable genes as input features for clustering analysis and downstream annotation. The Seurat packages FindNeighbors and FindClusters were used for clustering at a resolution of 0.5. Uniform manifold approximation and projection (UMAP) was utilized for visualization of clustering. For marker identification and annotation of clusters, Seurat’s FindMarkers tool was used with a requirement that the markers were expressed in 25% of the cells present in the dataset. For some rare population detection, the 25% criterion was removed. From the FindMarkers results, we consider genes with log2 fold changes that are greater than 2, and adjusted p-value less than 0.05 as significantly differentiated genes. If a cluster did not contain any genes that pass this threshold, we did not consider this as a significant cluster.

---

**Figure S1.**
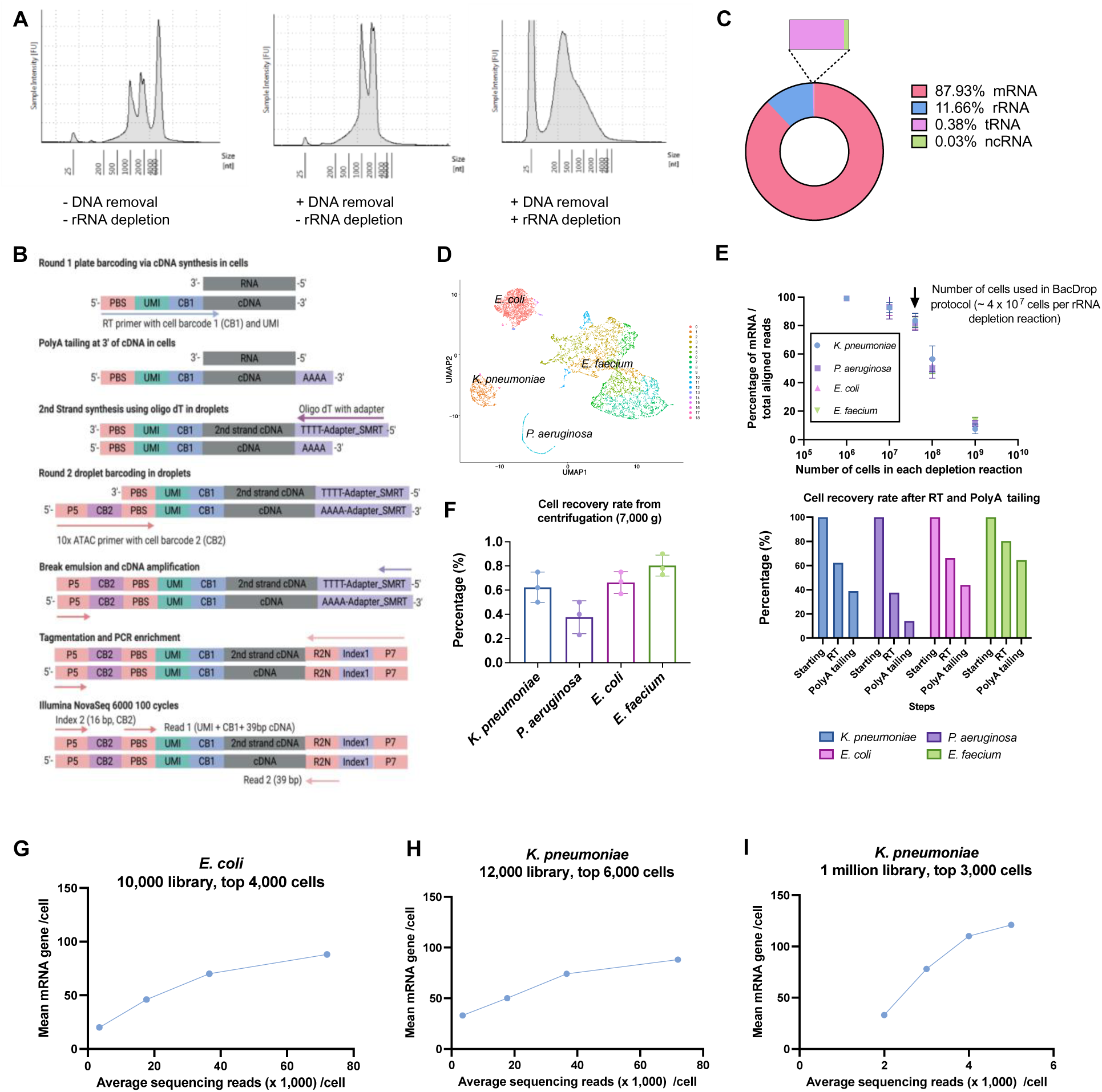
Development and performance of BacDrop. (A) rRNA and gDNA depletion are implemented in BacDrop. RNA was extracted from permeabilized cell with or without rRNA and gDNA depletion. The three peaks are 16s rRNA (∼1000 bp), 23 rRNA (∼2000 bp), and gDNA (> 4000 bp). **(B)** Scheme of two rounds of cell barcoding and library construction. **(C)** RNA species and their percentage in a BacDrop library of *K. pneumoniae*. **(D-F**) Validation of BacDrop in four bacterial species, including *K. pneumoniae*, *P. aeruginosa*, *E. coli*, and *E. faecium*. **(D)** Unsupervised cell clustering separates these four species into distinct clusters. Multiple subpopulations of *E. faecium* were identified due to a small differential expression (Log2 fold change 0.6 – 0.7) of two highly expressed housekeeping genes (*ef-tu* and *ef-g*). **(E)** rRNA depletion efficiency in these four species. Due to the capacity of the rRNA depletion kit (10 µg total RNA maximal), the rRNA depletion efficiency is a function of cell numbers. In BacDrop protocol, we used 4 x 10^7^ cell for each rRNA depletion reaction, resulting in a ∼80% depletion efficiency. **(F)** Cell loses in the mixed-species experiment are due to different cell recovery rates from centrifugation steps. In this experiment, equal numbers of cells from each species were mixed before round 1 plate barcoding (RT). Due to different cell sizes, these four species had different recovered rates at each centrifugation step at 7,000 *g*. After RT and PolyA tailing (right before round 2 droplet barcoding), the cells recovered from each species differed significantly. Therefore, we recommend users to optimize centrifugation speed for different species. In cases multiplexing is needed, we recommend keep samples separate until round 2 droplet barcoding. Experiments in (E and F) were repeated three times. Error bars are plotted as standard deviation. **(G-I)** Assessment of the transcriptome coverage of BacDrop libraries. **(G)** A library containing 10,000 cells of *E. coli* sequenced at 80,000 reads per cell. Roughly ∼4,000 cells were recovered with an average of 90 mRNA genes detected per cell. **(H)** A library containing 12,000 cells of *K. pneumoniae* sequenced at 80,000 reads per cell. Roughly ∼6,000 cells were recovered with an average of 88 mRNA genes detected per cell. **(I)** A library containing ∼1 million cells of *K. pneumoniae* sequenced at 5,000 reads per cell. Top 3,000 cells were analyzed with an average of 127 mRNA genes detected per cell. For libraries in (G and H), sequencing reads were randomly subsampled to ∼40,000, ∼20,000, ∼4,000 reads per cell, and the analysis was repeated to calculate numbers of mRNA genes detected per cell. For library in (I), sequencing reads were randomly subsampled to ∼4,000, ∼3,000, ∼2,000 reads per cell, and the analysis was repeated to calculate numbers of mRNA genes detected per cell.

**Figure S2.**
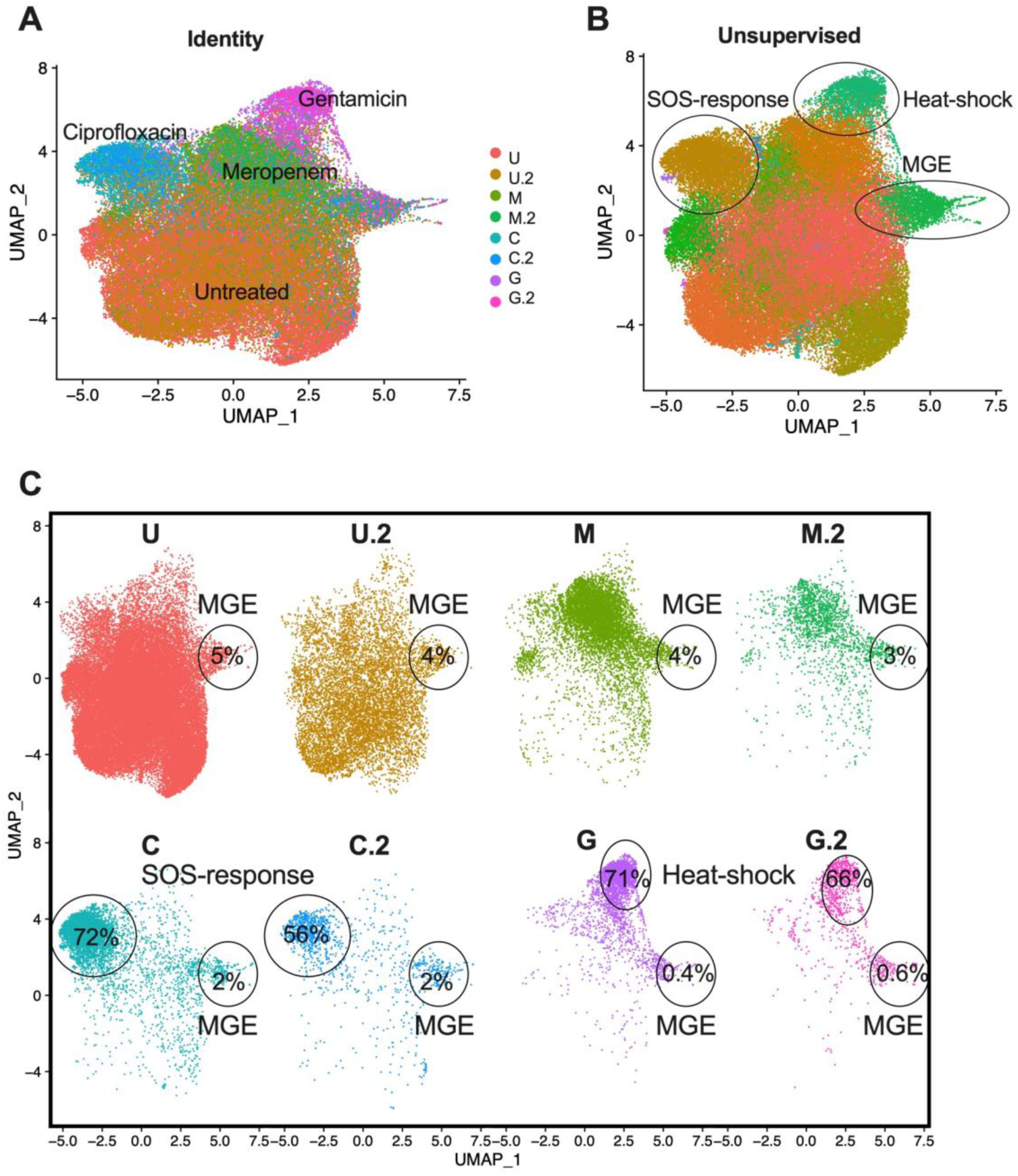
Analysis of MGH66 experiment including the antibiotic-treated samples and untreated samples. **(A)** UMAP plotted based on the original identity of these 8 samples. Cells were separated well based on their treatment. **(B)** Unsupervised UMAP showed three clusters with significantly (p < 0.05) higher expression of genes in the SOS-response pathway, heat-shock response, and genes encoding an IS903B transposase (MGE). **(C)** No strong batch effect was observed between the two biological replicates with the same treatment conditions. 56%-72% of the ciprofloxacin-treated cells were clustered in the SOS-response cluster, and 66%-71% of the gentamicin-treated cells were clustered in the heat-shock response cluster. All 8 samples contained cells highly expressing IS903B (MGE cluster).

**Figure S3.**
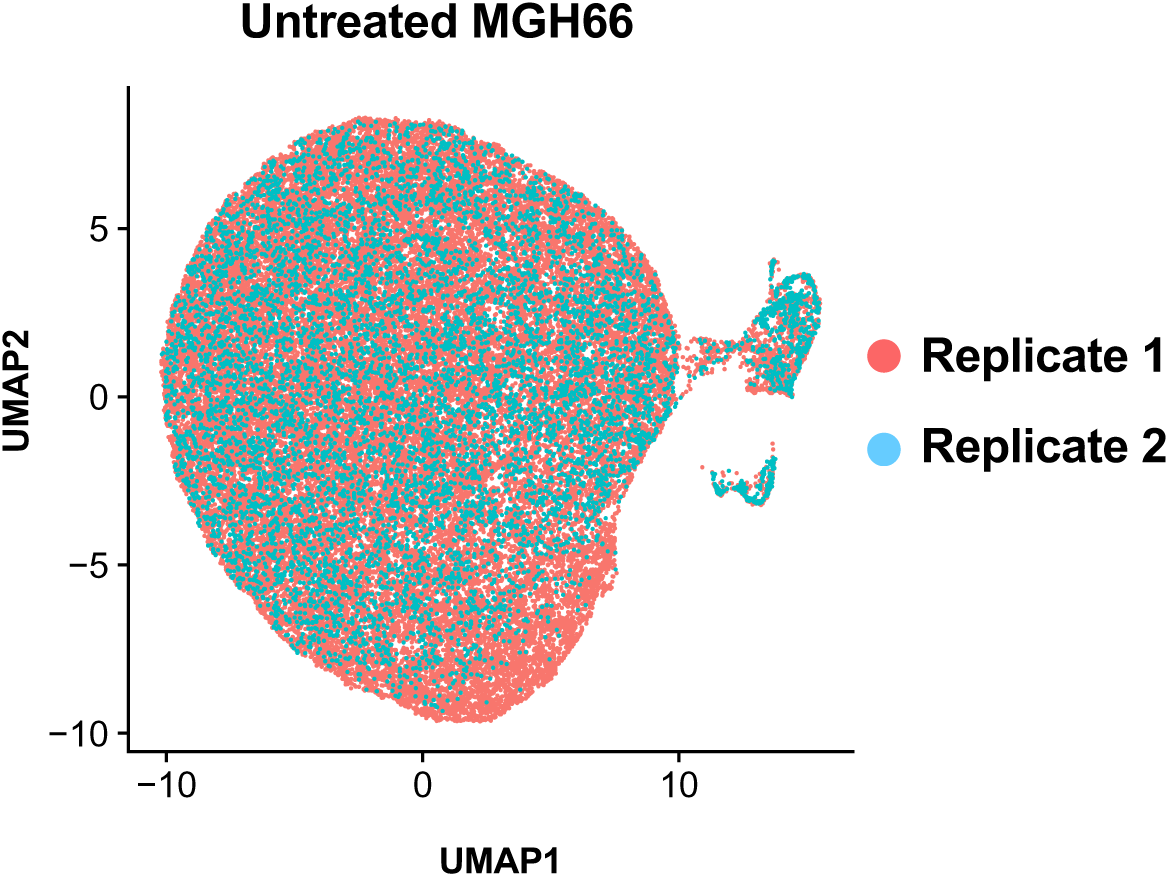
Analysis of the untreated MGH66 samples. UMAP plotted based on the original identity of these two samples (replicate 1 and replicate 2).

**Figure S4.**
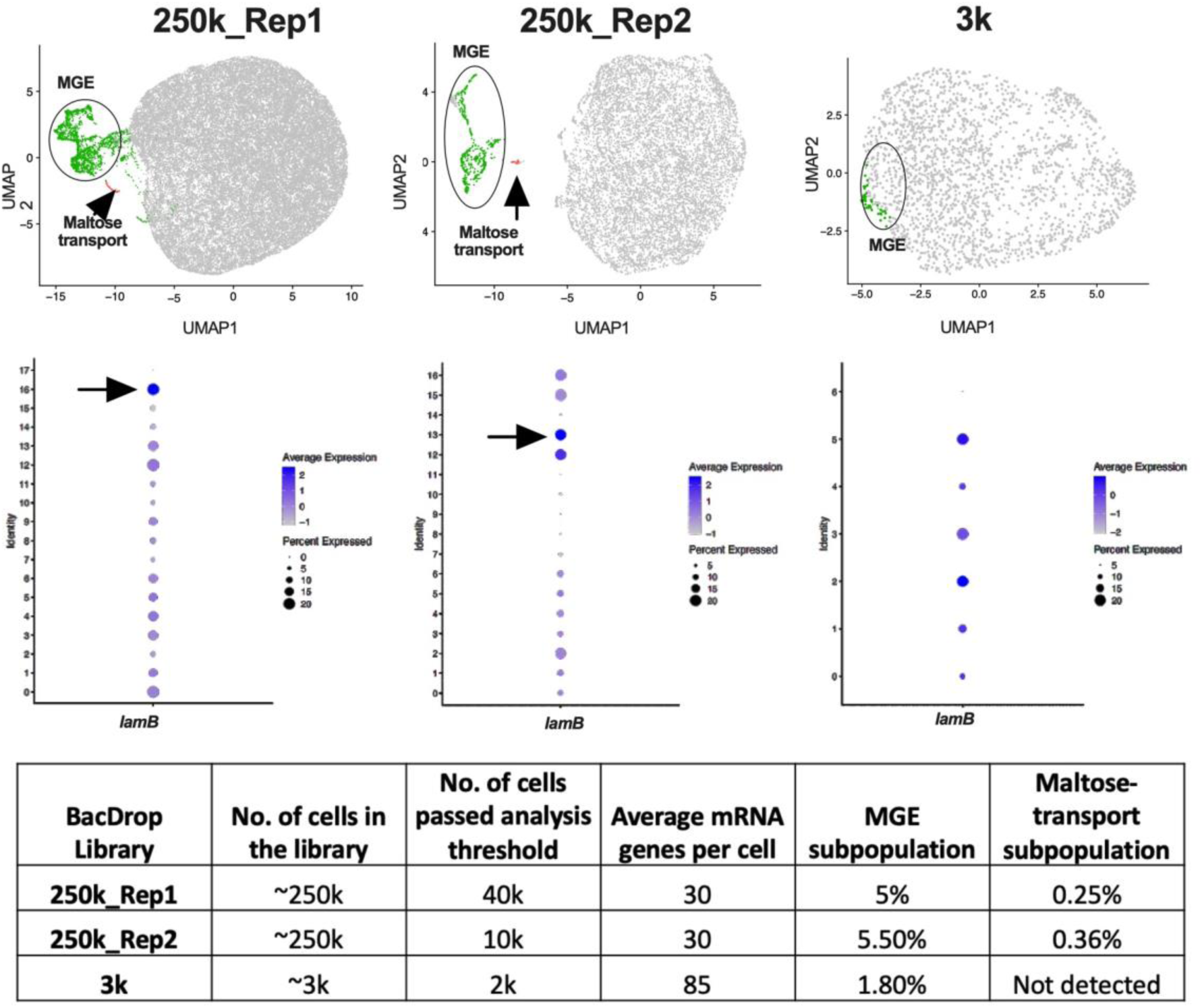
A large number of cells is beneficial for the detection of small population. Both libraries of Replicate 1 and Replicate 2 were constructed with ∼250k untreated MGH66 cells. We sequenced Replicate 1 with ∼5,000 reads/cell and Replicate 2 with ∼3,000 reads/cell, and recovered ∼40k cells and ∼10k cells from Replicate 1 and Replicate 2, respectively. In these two libraries, a subpopulation highly expressing maltose transport genes (i.e, *lamB*) were detected. But this subpopulation was not detected in the library containing only ∼3,000 untreated MGH66 cells that were collected at the same condition. The MGE subpopulation was detected in all libraries.

**Figure S5.**
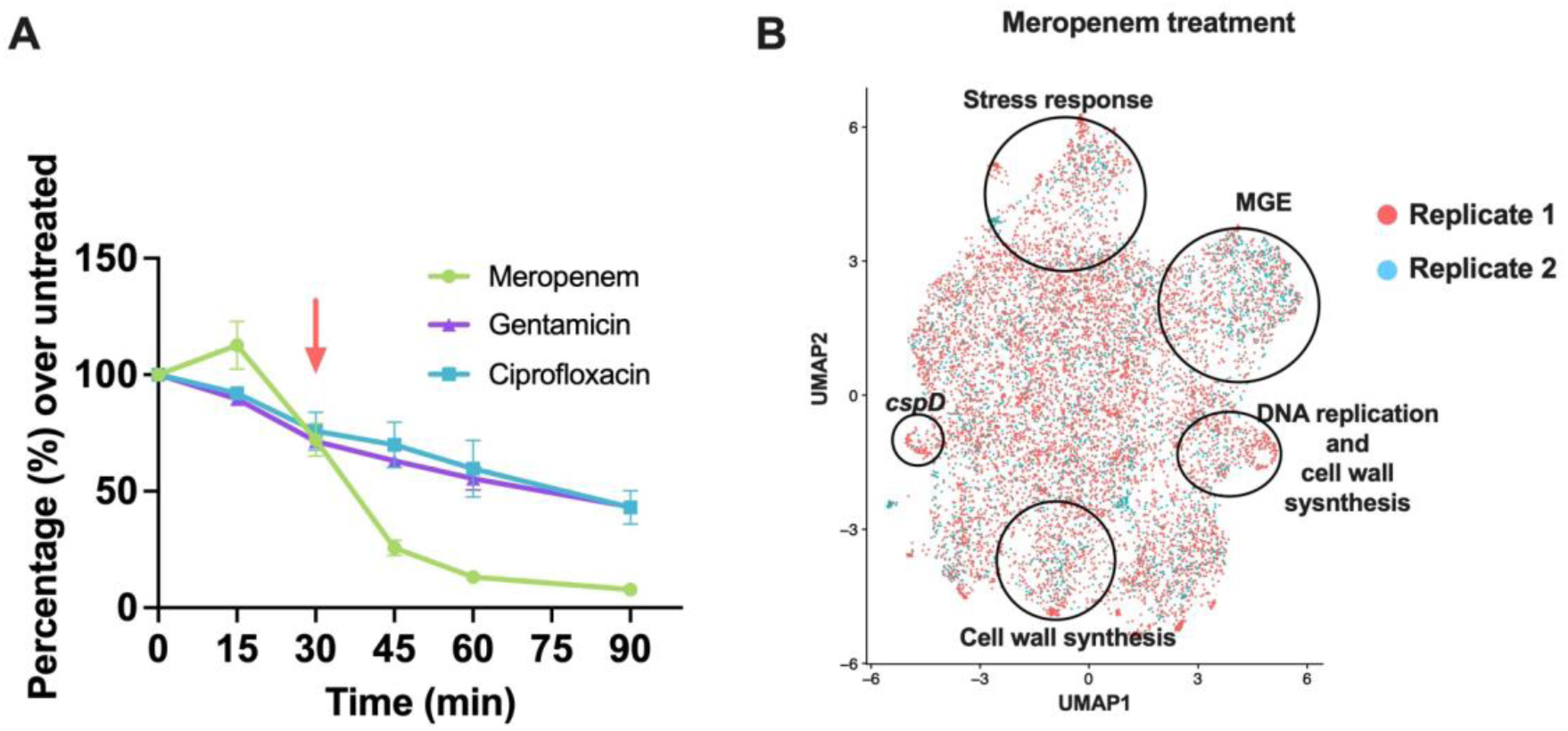
Analysis of the meropenem-treated samples. **(A)** Killing kinetics of MGH66 treated with antibiotics. At each time points, cells from meropenem treatment (green), Ciprofloxacin treatment (blue), Gentamicin treatment (purple), and the untreated samples were diluted and plated on LB agar plates without antibiotics. CFUs from each condition were normalized to the CFUs of the untreated samples at the same time points to calculate the killing rates for each time points. The experiment was repeated three times and the error bars are plotted as standard deviation. **(B)** Analysis of the meropenem-treated samples. UMAP plotted based on the original identity of these two samples (replicate 1 and replicate 2).

**Figure S6.**
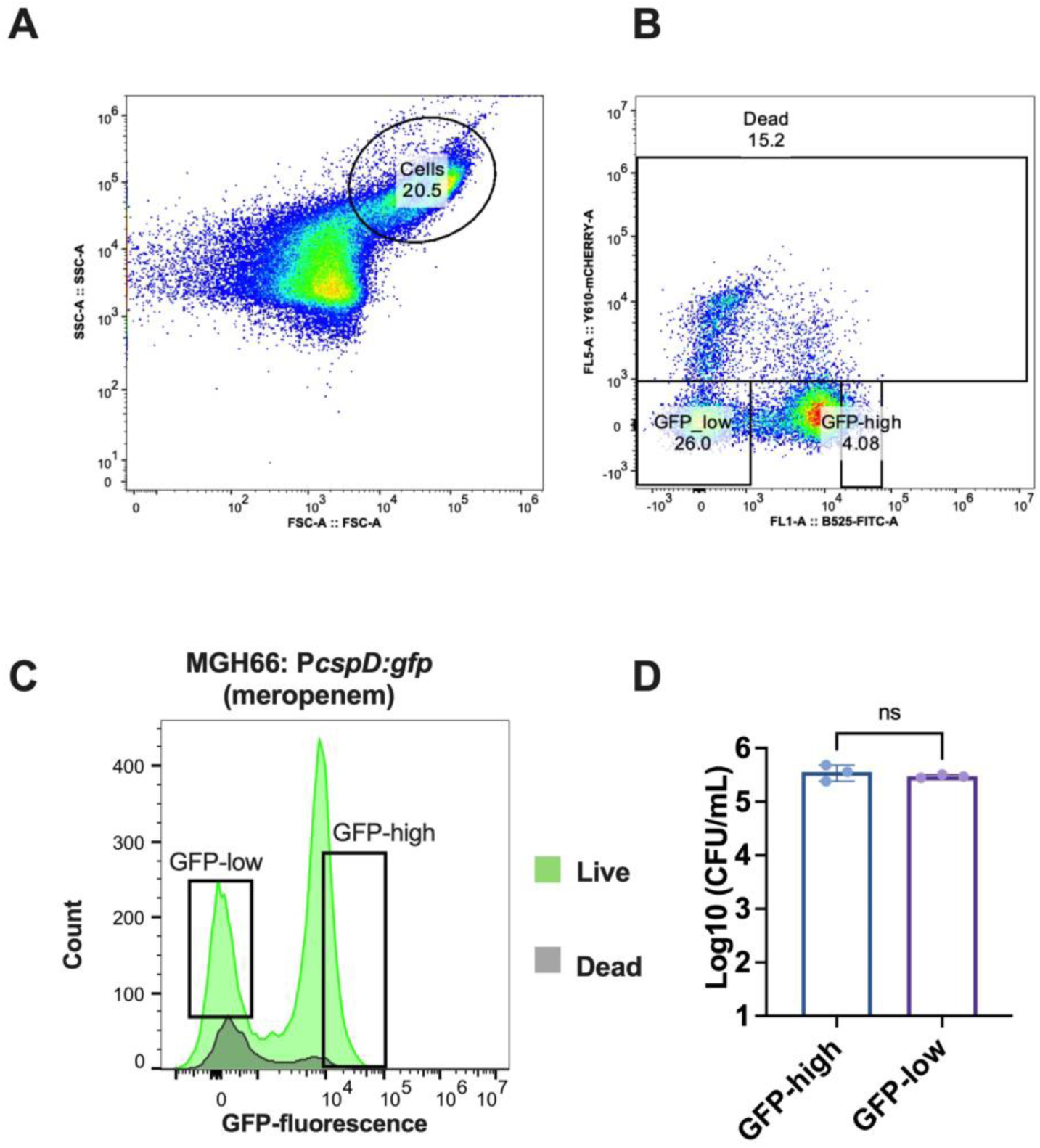
Validate the expression of *cspD* in MGH66: P*cspD*:gfp strain using FACS sorting coupled with Live/Dead staining. At 30 minuets of the meropenem treatment, cells were diluted into propidium iodide staining buffer (10^6^ cells/mL) and incubated at room temperature for 15 min in the dark. Then the cells were run through flowcytometry or subject to FACS sorting. **(A)** Gating for cells to exclude cell debris. **(B-C)** Dead cells were detected using mCherry red fluorescence, and GFP fluorescence of all cells was detected using green fluorescence. Roughly ∼15.2% cells are dead in the gated population. The population showing no mCherry fluorescence is live cells. The gating of GFP-low and GFP-high population is shown. **(D)** 10^4^ live cells from the GFP-high and GFP-low subpopulations were sorted into 1 mL LB medium, and immediate plated 100 µL on LB agar plates without any antibiotics. This experiment was run in triplicates and error bars were plotted as standard deviation. The student’s t-test was used for statistical analysis.

